# Muti-omics characterization reveals brain-wide disruption of synapses and region- and age-specific changes in neurons and glia in *Sp4* mutant mice, a genetic model of schizophrenia and bipolar disorder

**DOI:** 10.1101/2024.10.12.618006

**Authors:** Min Jee Kwon, Kira A. Perzel Mandell, Sameer Aryal, Chuhan Geng, Ally A. Nicolella, Antia Valle-Tojeiro, Sahana Natarajan, Deeksha Misri, Alyssa Hall, Bryan J. Song, Zohreh Farsi, Hasmik Keshishian, Steven A. Carr, Morgan Sheng

## Abstract

Schizophrenia and bipolar disorder are highly heritable mental illnesses with unclear pathophysiology. Heterozygous loss-of-function mutations of *Sp4*, a zinc-finger transcription factor, greatly increase risk of schizophrenia and bipolar disorder. To investigate the molecular functions of *Sp4* in an unbiased manner *in vivo*, we performed multi-omics analyses of *Sp4* mutant mice. Bulk and single nucleus RNA-seq data showed prominent gene expression changes in all brain regions and most cell types, including neuronal and non-neuronal cells. Gene set enrichment analysis of transcriptomic changes revealed alterations in many molecular pathways, including synapse, oxidative phosphorylation, and ribosome. Synapse proteomics of *Sp4* mutants pointed to impaired glutamatergic signaling and altered presynaptic function. In *Sp4* heterozygous mutant mice, prefrontal cortex and striatum exhibited downregulation of synapse pathways and neuronal hypoactivity at 1 month, associated with reduced sterol biosynthesis in astrocytes, whereas at 3 months, there was a shift to neuronal hyperactivity, concurrent with suppressed immune pathways in the striatal microglia. Furthermore, our study found that much of the transcriptomic changes might be accounted for by a set of transcription regulators (*Nr3c1*, *Creb1*, and *Kdm5b*) under the control of *Sp4*. Overall, this study provides cellular and molecular features resulting from *Sp4* LoF that may explain the pathophysiology of SCZ-BD psychotic disorder spectrum.

## Introduction

Schizophrenia (SCZ) and bipolar disorder (BD) are severe psychiatric disorders that together affect approximately 1∼4% of the global population^1–4^. Although these psychotic disorders are highly heritable (estimated heritability of 60-80%^5–7^), their underlying biology and pathophysiology remain largely elusive. Recently, exome sequencing studies of case-vs-control human populations have discovered a set of genes in which rare loss-of-function mutations (in heterozygous state) greatly increase the risk for SCZ^8^ and BD^9^. Identification of these rare genetic variants with large impact (odds ratio 2-60 for SCZ and 3-40 for BD) has facilitated the investigation of the neurobiological mechanisms of SCZ and BD, in part by enabling the creation of animal models with bona-fide human genetics validity^10–12^.

*Sp4* has been identified as a rare-variant large-effect risk gene for SCZ by SCHEMA (Schizophrenia Exome Sequencing Meta-analysis)^8^, as well as a common-variant locus for SCZ by GWAS (Genome-Wide Association Study)^13^. In addition, heterozygous loss-of-function (LoF) mutations in *Sp4* are associated with BD^14,15^, making *Sp4* particularly interesting as a shared risk gene for SCZ and BD. *Sp4* is a member of the Sp (specificity protein) transcription factor (TF) family that encodes a zinc-finger TF that binds to GC- and GT-rich regions in the promoters of a large number of genes. *Sp4* is predominantly expressed in the brain, especially in neurons, and plays crucial roles in neurodevelopment and in the maintenance of neuronal function^16^. For example, knock-down of *Sp4* in rat primary cultured neurons results in impaired activity-dependent dendrite patterning during neuronal morphogenesis^17,18^, and *Sp4* homozygous null mice have reduced brain size and altered cortical layering^19^. In addition, mice with a hypomorphic *Sp4* mutation have abnormal behaviors, such as prepulse inhibition defects that were reversed upon germline-cre-dependent restoration of *Sp4* levels^20^. Furthermore, reduced expression of *Sp4* has been reported in the brain regions including prefrontal cortex and hippocampus of SCZ and BD patients^21^ and in lymphocytes of patients with first-episode psychosis^22^.

Unlike some other SCZ risk genes, *Sp4* has not been associated with intellectual disability or developmental disorders but rather seems to be a relatively specific risk gene for the SCZ-BD psychotic disorder spectrum^8^. Despite its importance, *Sp4* has not been extensively studied in the brain. In this study, we used unbiased comprehensive multi-omic approaches to explore in detail the consequences of heterozygous *Sp4* LoF across multiple brain regions and cell types, within cortical synaptic compartments, and at two different postnatal ages. Our data reveal that *Sp4* LoF has an extensive impact on neuronal gene expression throughout the brain, likely via regulation of a set of other TFs and chromatin modifiers that are critical for neuronal function and signaling. This study provides a molecular-to-systems-level picture of the *Sp4* mutant brain that facilitates deeper understanding on the pathophysiology and disease mechanisms of SCZ and BD.

## Results

### Large-scale widespread changes in brain transcriptome in *Sp4* mutant mice

We used CRISPR/Cas9 technique to generate *Sp4* LoF mutant mice with an early terminating stop codon in Exon3 (Figure 1A and B), where *Sp4* genetic variants are concentrated in SCZ patients^8^. We confirmed that the Sp4 protein levels in brains were reduced in heterozygous (*Sp4+/-*) and undetectable in homozygous (*Sp4-/-*) mutant mice (Figure 1C). Previously, homozygous *Sp4* mutant mice (using various knockout strategies) were reported to show growth retardation, developmental problems, and peri- and post-natal lethality^16,23,24^. Our CRISPR/Cas9-generated homozygous *Sp4* mutant mice consistently showed peri- and post-natal lethality and a reduction in body and brain weights in surviving males and females at 1- and 3-months of age (Figure 1D and E; Figure S1A and B). *Sp4+/-* mutants had similar body and brain weights compared to their wild-type (WT) littermates (Figure 1D and E; Figure S1A and B).

**Figure 1.**
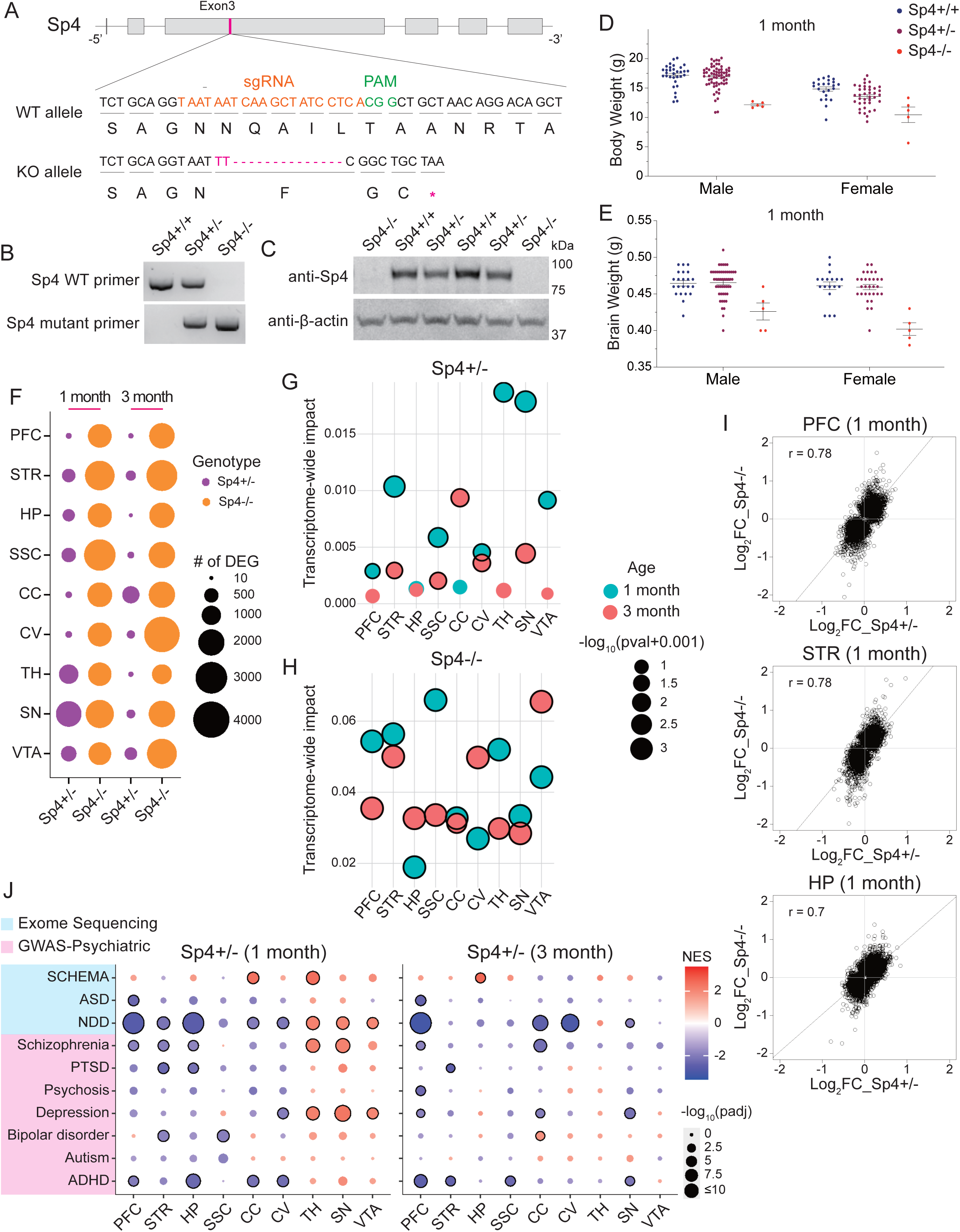
Brain region- and age-specific transcriptomic changes in *Sp4* mutant mice. (A) Schematic diagram of *Sp4* mutant mouse generation using CRISPR-Cas9 technique. Exon3 of *Sp4* is targeted by sgRNA to have an early stop codon in the mutant mice. (B) Genotyping results of *Sp4+/+*, *Sp4+/-*, and *Sp4-/-* mice using WT and mutant primers of *Sp4*. (C) Western blots probing for SP4 and β-actin in total brain lysate. (D and E) Body (D) and brain (E) weights of the 1-month-old *Sp4+/-* and *Sp4-/-* mutants compared to WT littermates. (F) Number of differentially expressed genes (DEGs) in the indicated brain regions and ages in *Sp4+/-* and *Sp4-/-* mutants. (G and H) Transcriptome-wide impact (TWI) scores analyzed across the indicated brain regions and ages for *Sp4+/-* (G) and *Sp4-/-* (H) mutant mice. (I) Gene expression correlation in the indicated brain regions and genotypes at 1 month. Spearman’s r correlation values are indicated on the plots. Log_2_FC, log_2_Fold Change. (J) GSEA results for risk genes associated with psychiatric and neurological diseases. Circles with black outlines indicate statistical significance (adjusted p-value < 0.05). NES, normalized enrichment score. In (D) and (E), each circle represents an individual animal. In (G) and (H), bubble size indicates -log_10_(pval+0.001), color indicates age, and circles with black outlines indicate statistical significance (p-value < 0.05). In (J), ASD, autism spectrum disorder; NDD, neurodevelopmental disorder; PTSD, post-traumatic stress disorder; ADHD, attention deficit hyperactivity disorder. See also Figure S1.

To determine the effect of *Sp4* LoF on gene expression in the brain, we first performed bulk RNA sequencing (RNA-seq) analysis on *Sp4+/-*, *Sp4-/-*, and WT littermates at 1 and 3 months of age across multiple brain regions, including the prefrontal cortex (PFC), hippocampus (HP), somatosensory cortex (SSC), striatum (STR), thalamus (TH), substantia nigra (SN), ventral tegmental area (VTA), cerebellar cortex (CC), and cerebellar vermis (CV). Samples from the same brain regions clustered together in the principal component analysis (PCA), implying the consistency of tissue dissection (Figure S1C-F). According to bulk RNA-seq, *Sp4* was widely expressed across all examined brain regions and most highly in cerebellum (both CC and CV) as previously reported^25,26^ (Figure S1G). *Sp4* expression was higher in cerebellum (CC and CV), PFC, SSC, and TH at 3 months compared to 1 month in WT animals (Figure S1G).

Differential expression (DE) analysis of the bulk RNA-seq data identified numerous differentially expressed genes (DEGs; padj < 0.05) in *Sp4+/-* and *Sp4-/-* mutants compared to WT littermates in all nine brain regions and at the two ages studied (Figure 1F). *Sp4-/-* mutants exhibited widespread and particularly large changes in their brain transcriptome (1067-3825 DEGs in most brain regions at both 1 and 3 months) (Figure 1F). Although generally less impacted than homozygous mutants in terms of DEGs, *Sp4+/-* mice nonetheless showed major transcriptomic changes across the brain, especially in SN, TH, and VTA at 1 month (1887, 1016, and 606 DEGs, respectively) and CC, VTA, and STR at 3 months (774, 375, and 216 DEGs, respectively) (Figure 1F). In keeping with DEG counts, transcriptome-wide analysis of differential expression (TRADE)^27^ showed that *Sp4+/-* mutants generally had greater transcriptomic changes at 1 month than at 3 months in most brain regions (Figure 1F and G), whereas for *Sp4-/-* mutants, the transcriptomic changes were similarly large at 1 and 3 months of age (Figure 1F and H).

Consistent with *Sp4-/-* having more DEGs, correlation plots of Log_2_ fold change (Log_2_FC) of the unions of nominally significant genes (p-value < 0.05) between heterozygous and homozygous *Sp4* mutants showed that *Sp4-/-* mutants on average had larger Log_2_FCs than *Sp4+/-* mutants in most brain regions at both 1 and 3 months (Figure 1I; S1H and I). Yet, transcriptome-wide changes in *Sp4+/-* and *Sp4-/-* mutants were generally highly correlated in all nine brain regions at both 1 and 3 months (Spearman’s r = 0.6–0.85; except for CV at 3 months where Spearman’s r = 0.33; Figure 1I; Figure S1H and I). Together, these analyses indicate that loss of one copy of *Sp4* is sufficient to result in large-scale changes of gene expression in the brain that are considerably overlapping with, though less severe than, those found in the homozygous *Sp4* LoF mutant. Given that SCHEMA gene mutations are associated with human SCZ in heterozygous state, and because homozygous mutant mice have obvious developmental abnormalities (Figure 1D and E; Figure S1A and B), we focused our subsequent analyses on *Sp4+/-* mutants, which are the more appropriate model for SCZ/BD.

Are the genes associated with psychiatric and neurological disorders^28–30^ (Table S1) enriched among the differentially expressed genes in *Sp4+/-* mutants? By gene set enrichment analysis (GSEA)^31,32^, we found that genes associated with SCZ (by GWAS) were significantly enriched among downregulated genes in PFC, STR, and HP, and among upregulated genes in TH and SN, in 1-month-old *Sp4+/-* mutants (Figure 1J). Neurodevelopmental disorder (NDD)-associated genes showed a similar pattern of gene-set enrichment as schizophrenia-associated genes, in keeping with the known overlap in genetic risks between SCZ and NDD^8,33^ (Figure 1J). Considering that *Sp4* is linked to BD, it was interesting that genes associated with depression were enriched in upregulated genes of TH, SN, and VTA in *Sp4+/-* mutants (Figure 1J). GSEA signals for SCZ or NDD genes were generally diminished at 3 months of age, except for PFC and CC (Figure 1J) Thus, *Sp4* heterozygous LoF alters expression of multiple sets of genes that are genetically linked to SCZ and related psychiatric disorders.

### Region-specific changes in synapse and other molecular pathways in *Sp4* mutants

Based on the GSEA analysis of bulk RNA-seq data, we found that *Sp4* LoF resulted in dysregulations in multiple pathways (annotated by gene ontology [GO] terms) across widespread brain regions. Prominent changes occurred in GO terms related to synapses, ribosomes, oxidative phosphorylation, neuron development, and RNA processing/splicing that have been consistently dysregulated in the previous studies of human brain tissues and animal models of SCZ^10,34–36^ (Figure 2A; Figure S2A). In both *Sp4+/-* and *Sp4-/-* mutants, synapse-related GO terms, including glutamatergic and GABAergic synapses, were downregulated in most of the brain regions examined (PFC, STR, HP, SSC, CC, and CV) but upregulated in TH, SN, and VTA at 1 month (Figure 2A; Figure S2A). GSEA with more refined synaptic terms (SynGO)^37^ showed alterations in synaptic genes with largely the same region-specific pattern as the standard GSEA (Figure 2B, compare with Figure 2A). SynGO analysis also revealed that both presynaptic and postsynaptic processes were strongly affected in *Sp4+/-* mutants at 1 and 3 months (Figure 2B). Interestingly, “Translation at Presynapse” and “Translation at Postsynapse” SynGO terms were significantly upregulated in PFC, HP, CC, and CV but downregulated in SSC, TH, SN, and VTA in *Sp4+/-* mutants at 1 month, which is the opposite directional change as the other presynapse- and postsynapse-related GO terms (Figure 2B; see also Figure 2A “Ribosome” GO term). Thus, changes in translation-related gene expression are anti-correlated with changes in other synapse pathways (Figure 2B). Similar inverse relationship between translation-/ribosome-related GO terms and synapse-related GO terms were found in *Grin2a* mutant mice, another genetic animal model of SCZ^10^.

**Figure 2.**
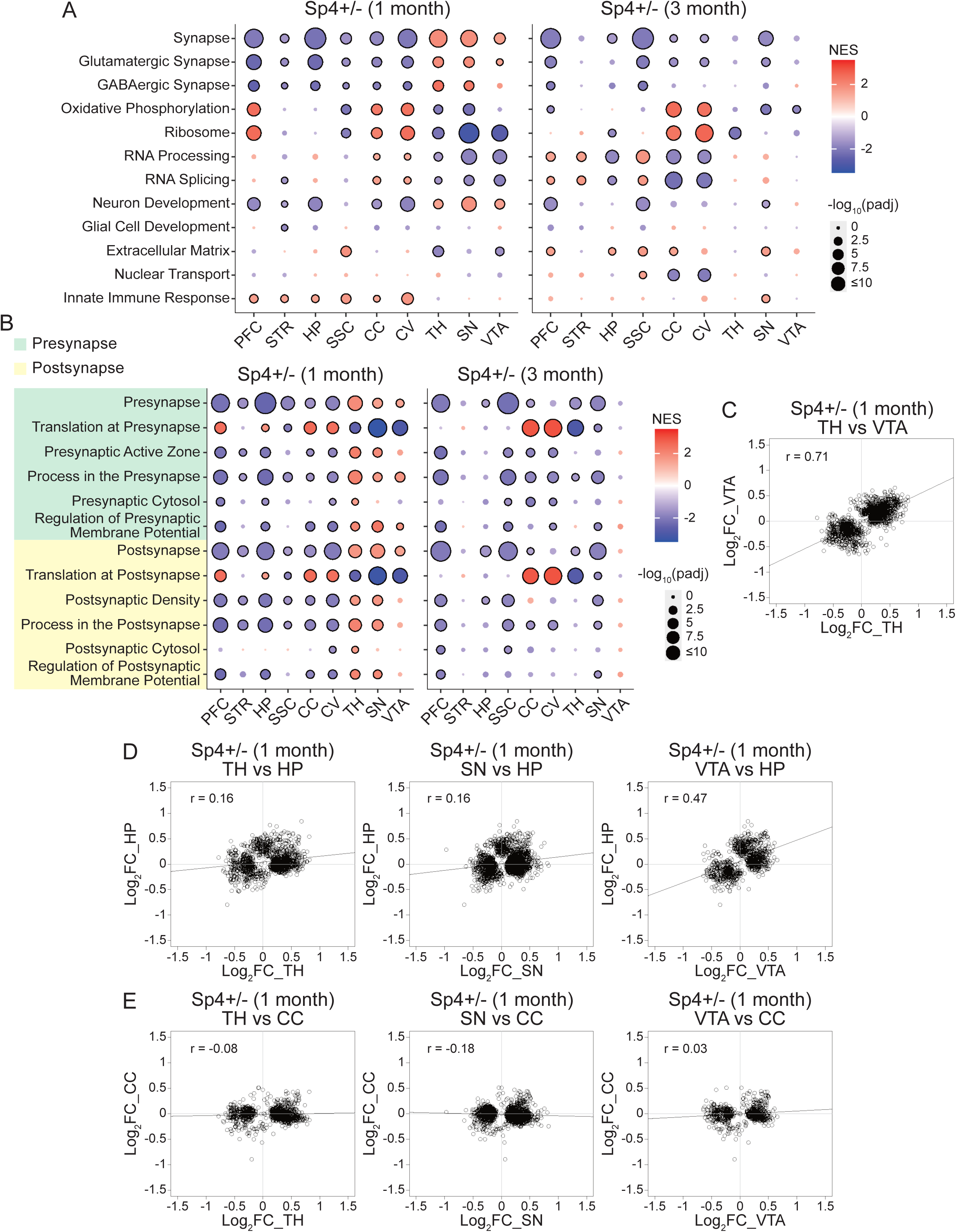
Transcriptomic changes in multiple molecular pathways in *Sp4* mutant mice. (A and B) GSEA results of *Sp4+/-* mutants in the indicated brain regions and ages for a selection of GO terms from MSigDB (A) and SynGO (B). (C) Correlation plot comparing the Log_2_FC of the DEGs between TH and VTA of *Sp4+/-* mutants at 1 month. (D) Correlation plots comparing the Log_2_FC of the DEGs between TH, SN, or VTA versus HP of *Sp4+/-* mutants at 1 month. (E) Correlation plots comparing the Log_2_FC of the DEGs between TH, SN, or VTA versus CC of *Sp4+/-* mutants at 1 month. In (A) and (B), circles with black outlines indicate statistical significance (adjusted p-value < 0.05). NES, normalized enrichment score. In (C-E), Spearman’s r correlation values are indicated on the plots. Log_2_FC, log_2_Fold Change. See also Figure S2.

What specific genes are dysregulated in *Sp4* mutant brains? In line with GSEA results of psychiatric disorder gene sets (see Figure 1J), several SCZ risk genes from SCHEMA such as *Akap11*, *Xpo7*, *Nr3c2*, and *Hcn4,* and members of the *Mef2* family of TFs (*Mef2a*, *Mef2c*, and *Mef2d*) that are known to be associated with various neurodevelopmental and psychiatric disorders, including SCZ^38–40^, were significantly increased in TH of *Sp4+/-* mutants at 1 month (Figure S2B). Additionally, the expression of many neurotransmitter receptors and signaling molecules such as *Gabra1*, *Gabrb1*, *Gabrb2*, *Gabrg3*, *Gria4*, *Npy1r*, and *Gsk3b* were significantly elevated in TH of 1-month-old *Sp4+/-* mutants (Figure S2B). However, these significant changes observed in 1-month-old TH of *Sp4+/-* mutants were largely lost and even reversed at 3 months of age (data not shown).

Also, consistent with the GSEA results (see Figure 2A and B), many of the down-regulated DEGs in STR can be described in the context of synaptic function and signaling, such as *Pclo*, *Reln*, *Grid1*, *Nbea* (at 1 month, Figure S2C), *Nptxr*, *Maob* (at 3 months, Figure S2D), *Cd59a*, and *Ryr1* (both 1 and 3 months, Figure S2C and D). Intriguingly, unlike TH, many DEGs of STR in *Sp4+/-* mutants were overlapping between 1 and 3 months (Figure S2C and D), indicating convergent and prolonged transcriptomic changes in STR resulting from heterozygous LoF of *Sp4*.

In the GSEA analysis of the nine brain regions, we noticed a striking segregation between TH, SN, and VTA versus the other six brain regions examined (PFC, STR, HP, SSC, CC, and CV) in both *Sp4+/-* and *Sp4-/-* mutants at 1 month of age. For instance, synapse-related GO terms were downregulated in PFC, STR, HP, SSC, CC, and CV but upregulated in TH, SN, and VTA (Figure 2A and B; Figure S2A). As another example, “Innate Immune Response” GO term was significantly upregulated in PFC, STR, HP, SSC, CC, and CV but showed no change in TH, SN, and VTA of *Sp4+/-* mutants at 1 month (Figure 2A). Notably, this contrasting pattern of changes in TH, SN, and VTA versus the other six brain regions was largely lost at 3 months (Figure 2A and B; Figure S2A). When we performed correlation analysis of the Log_2_FCs of DEGs among TH, SN, and VTA of *Sp4+/-* mutants, we found a high correlation of gene expression changes within these three regions at 1 month (Spearman’s r = 0.69-0.74; Figure 2C; Figure S2E and F). On the other hand, there was weak correlation between the DEGs of TH and SN with those of PFC and HP (Spearman’s r = 0.09-0.16) (Figure 2D; Figure S2G and H). Notably, there was little or even inverse correlation (Spearman’s r = -0.28-0.03) between the DEGs from TH, SN and VTA versus cerebellum (CC and CV) (Figure 2E; Figure S2J-L). It is intriguing that mutation of *Sp4*, a transcription factor that presumably acts primarily in a cell autonomous manner, would have opposite-direction effects on synaptic and other molecular pathways, depending on the brain region. This implies a functional effect of *Sp4* LoF at the level of brain circuits, occurring particularly during a juvenile period of brain maturation (1 month of age) when SCZ pathomechanisms are most relevant.

### Altered glutamatergic signaling and synaptic vesicle pathways in the synaptic proteome of *Sp4* mutants

Several lines of evidence including human genetics, point to synaptic dysfunction in SCZ^8,11,28^, and consistently, our transcriptomic analysis showed dysregulation of synapse-related genes and pathways (see above, Figure 2). To study the effects of *Sp4* LoF on synapses at the protein level, we performed quantitative mass spectrometry proteomics on synaptic fractions (PSD fractions, see Methods) purified from the cerebral cortex of 1- and 3-month-old *Sp4+/-* mutants and their WT littermates. Many interesting differentially expressed proteins (DEPs, see Methods) were revealed in the synapse fractions of the cortex of *Sp4+/-* mutants (Figure 3A and B).

**Figure 3.**
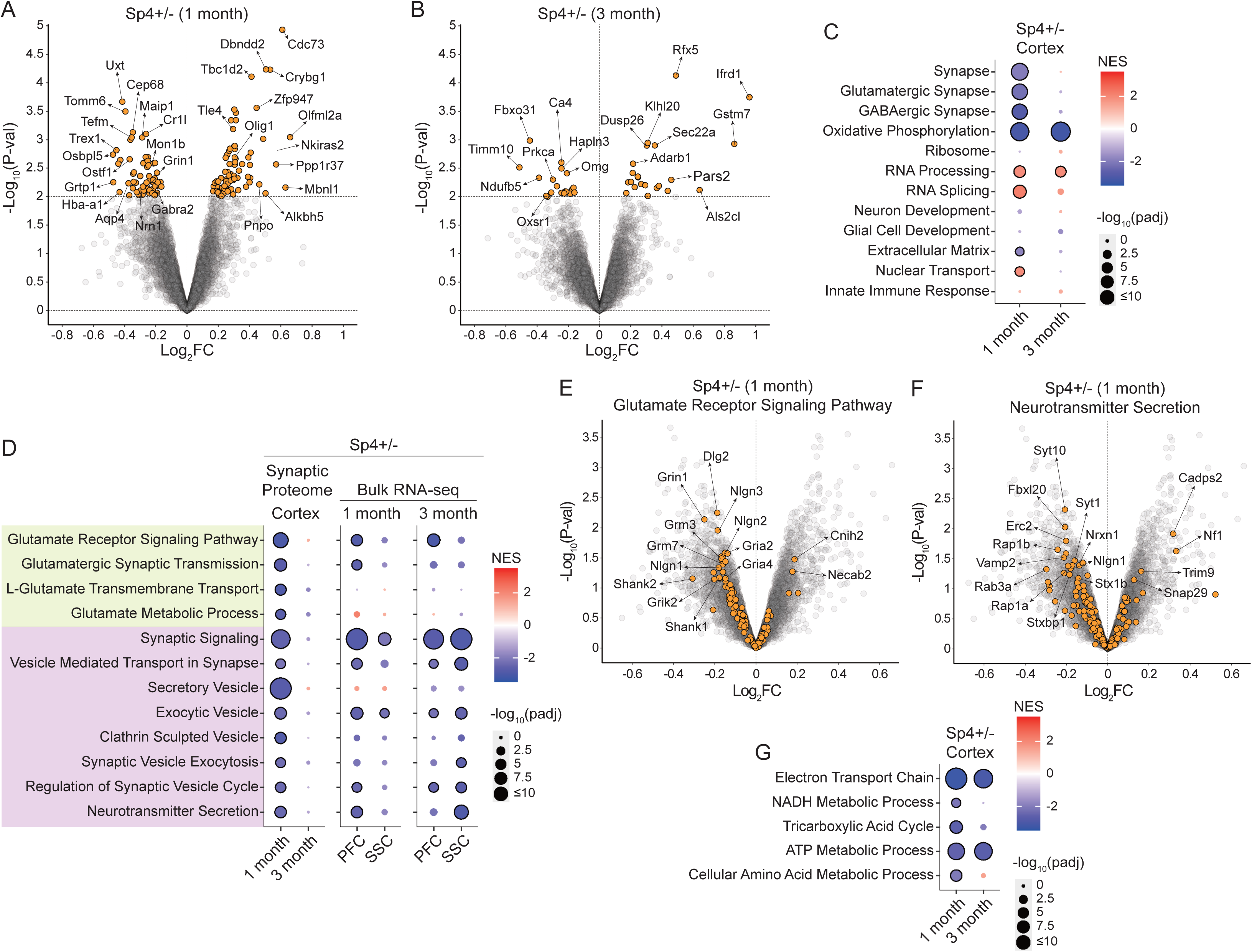
Decreased glutamate receptor signaling and synaptic vesicle-related signaling in synapse proteome of *Sp4* mutant mice. (A and B) Volcano plot of proteomic changes in the cortical synapses of *Sp4+/-* mutant mice at 1 (A) and 3 (B) months of age. (C) GSEA results of the cortical synapses of *Sp4+/-* mutants at the indicated ages for a selection of GO terms from MSigDB. (D) GSEA results for the indicated glutamate- and synaptic vesicle-related GO terms from the cortical synapse proteome and transcriptome of the indicated brain regions of *Sp4+/-* mutant mice at the indicated ages. (E and F) Volcano plots of proteomic changes in the cortical synapses of 1-month-old *Sp4+/-* mutant mice, highlighting genes from the “Glutamate Receptor Signaling Pathway” (E) and “Neurotransmitter Secretion” (F) GO terms. Log_2_FC, log_2_Fold Change. (G) GSEA results for the indicated energy metabolism-related GO terms in the cortical synapses of *Sp4+/-* mutant mice at the indicated ages. In (A) and (B), yellow dots indicate differentially expressed proteins (DEPs). In (C), (D), and (G), circles with black outlines indicate statistical significance (adjusted p-value < 0.05). NES, normalized enrichment score.

Among the downregulated DEPs in 1-month-old *Sp4+/-* synaptic fractions were: GRIN1, a constitutive subunit of the NMDA receptor that is crucial for synaptic plasticity and memory function^41^, and hypofunction of which is long implicated in SCZ; GABRA2, a subunit of the GABA-A receptor that is important for inhibitory signaling^42^; and NRN1, a glycophosphatidylinositol-linked protein that stimulates axonal plasticity, dendritic arborization and synapse maturation^43^ (Figure 3A). Furthermore, PRKCA (protein kinase C alpha), which is involved in various signaling pathways important for neural plasticity, learning and memory^44^, and OXSR1, a serine/threonine-protein kinase involved in response to oxidative stress and neuronal survival^45^ were significantly decreased in the synaptic proteome of *Sp4+/-* cortex at 3 months (Figure 3B). Among upregulated DEPs of *Sp4+/-* cortex at 3 months were ADARB1, an enzyme important for editing of glutamate receptor pre-mRNAs, neural signaling, and plasticity^46^, and IFRD1, an interferon-related protein involved in stress, immune responses, and inflammation^47,48^ (Figure 3B). A greater number of DEPs in synaptic fractions was found in *Sp4+/-* mutants at 1 month (131 DEPs) than at 3 months (41 DEPs) (Figure 3A and B), which is consistent with the greater transcriptomic impact in *Sp4+/-* mutants at 1 month (see Figure 1F and G). Thus, both transcriptomics and proteomics data indicate that the changes in the brain *of Sp4+/-* mutants subside considerably from 1 month to 3 months of age.

GSEA of the synapse proteomics data also showed more extensive pathway changes at 1 month in *Sp4+/-* mutants, affecting GO terms such as glutamatergic and GABAergic synapses, RNA processing, extracellular matrix and nuclear transport (Figure 3C). Many glutamate-related GO terms were significantly downregulated in the synapse of 1-month-old *Sp4+/-* cortex (Figure 3D), driven in large part by reduction of multiple glutamate receptor subunits (GRIN1, GRIK2, GRM3, GRM7, GRIA2, and GRIA4) and excitatory postsynaptic scaffolding proteins (DLG2, SHANK1, and SHANK2) (Figure 3E). Synaptic vesicle-related GO terms were also strongly downregulated in the synapse proteomics of *Sp4+/-* mutants at 1 month (Figure 3D), with a number of specific proteins involved in synaptic vesicle dynamics and neurotransmitter release (VAMP2, RAB3A, SYT1, SYT10, STX1B, and STXBP1) showing a decreased abundance (Figure 3F). These synapse proteomic changes in glutamate receptor- and synaptic vesicle-related pathways, which were concordant with bulk RNA-seq changes of the cortical regions PFC and SSC (Figure 3D), point to both pre- and postsynaptic disturbance in *Sp4+/-* mutant brain. Of note, the GSEA changes in glutamate receptor signaling and synaptic vesicle pathways were essentially gone at 3 months of age (Fig 3D), implying compensation and/or adaptation of pre- and postsynaptic processes in *Sp4+/-* mutants during brain maturation between 1 and 3 months. In contrast, we noted that GO terms related to mitochondria and respiration (such as oxidative phosphorylation, electron transport chain, and ATP metabolic process) were significantly downregulated in the synaptic proteome at both 1 and 3 months (Figure 3C and G). These findings suggest that synaptic energy metabolism is perturbed alongside excitatory neurotransmission in *Sp4+/-* cortex.

The reduction of multiple glutamate receptor proteins, especially NMDA receptor core subunit GRIN1 (Figure 3A and E), offers biochemical support for the long-standing glutamate/NMDA hypofunction hypothesis of SCZ pathophysiology^49^. In addition, it is interesting to find in the *Sp4+/-* synapse proteomics reduced levels of *Nlgn1*, *Nlgn2*, *Nlgn3*, and *Nrxn1*, which are synaptic adhesion molecules that have been genetically linked to SCZ^50–55^ (Figure 3E and F). In summary, these proteomics findings underscore that presynaptic as well as postsynaptic functions are likely affected in *Sp4+/-* brain.

### Large-scale transcriptomic changes in neuronal and non-neuronal cells in *Sp4* mutants

We conducted single nucleus RNA-seq (snRNA-seq) to determine which and how specific cell types are altered by *Sp4* heterozygous LoF, focusing on PFC and STR, brain regions that show robust changes by bulk RNA-seq and that are implicated in SCZ pathophysiology^56,57^. Observed cell clusters were annotated to different cell types (Figure 4A and B; Figure S3A and B), based on the expression of marker genes (Figure S3C-F). There was no significant difference in cell type proportions between *Sp4+/-* mutant and WT mice in PFC and STR at either 1 or 3 months of age (Figure S3G and H). DE analysis of snRNA-seq data (using the pseudocell algorithm, a mixed model approach^58^, see Methods) showed that the transcriptome of neurons (excitatory and inhibitory) and non-neuronal cells (astrocytes, microglia, oligodendrocytes, and OPC) of PFC and STR were highly affected in *Sp4+/-* mutants at 1 and 3 months (Figure 4C and D). Neuronal cell types such as pyramidal neurons of PFC layer 2/3 (L2/3IT) and layer 5/6 (L5IT and L6CT), and inhibitory spiny projection neurons (SPNs) of STR, were especially highly impacted, showing DEGs that numbered from hundreds to more than one thousand (Figure 4C and D). Among non-neuronal cell types, astrocytes (in PFC and STR) and oligodendrocytes (in STR) showed a sizable number of DEGs in *Sp4+/-* mutants (Figures 4C and D). Since *Sp4* is reported to be primarily expressed in neuronal cells and minimally expressed in glial cells^16,26^, we hypothesize that the considerable changes in astrocytes and oligodendrocytes are secondary to the effects of *Sp4* LoF in neurons.

**Figure 4.**
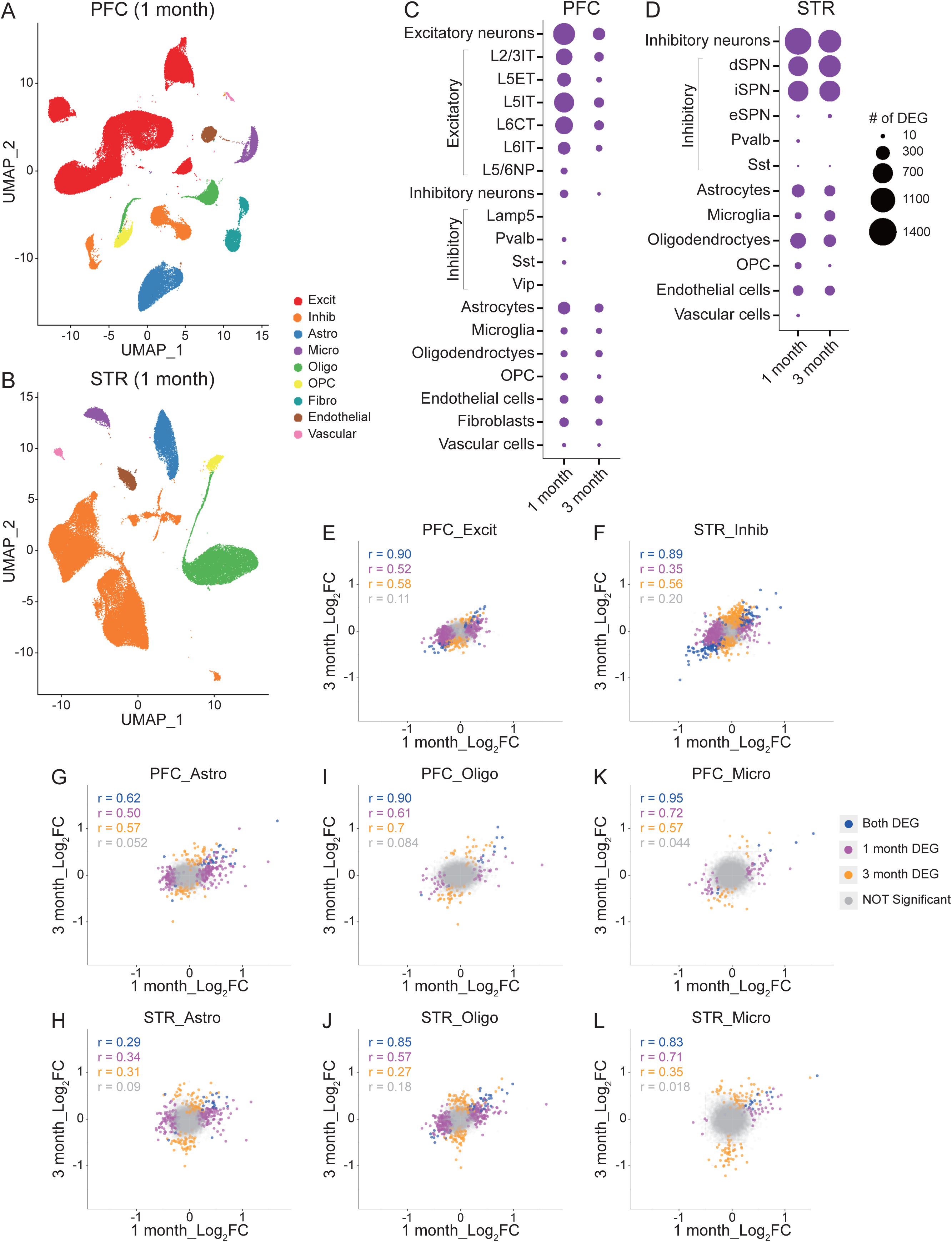
Cell type-specific transcriptomic changes in *Sp4* mutant mice. (A-B) Uniform manifold approximation and projection (UMAP) representation of the major cell types identified by snRNA-seq in the indicated brain regions at 1 month. (C-D) Number of DEGs across different cell types in the indicated brain regions and ages of *Sp4+/-* mutant mice. (E-L) Correlation plots of DEGs in the indicated cell types and ages of *Sp4+/-* mutant mice. Spearman’s r correlation values are indicated on the plots. Log_2_FC, log_2_Fold Change. In (A-D), OPC, oligodendrocyte progenitor cells; Pvalb, parvalbumin interneurons; Sst, somatostatin interneurons; Vip, vasoactive intestinal peptide interneurons; L2-L6, layers 2-6; IT, intratelencephalic; NP, near-projecting; ET, extratelencephalic; CT, corticothalamic neuron; dSPN, direct-pathway spiny projection neuron; iSPN, indirect-pathway SPN; eSPN, eccentric SPN. See also Figures S3.

Most of the cell types in STR, whether neuronal or non-neuronal, had similar number of DEGs at 1 and 3 months (Figure 4D), while PFC cell types generally had fewer DEGs at 3 months compared to 1 month (Figure 4C). Nonetheless, comparison of the log_2_FCs of DEGs between 1- and 3-month-old *Sp4+/-* mutants showed that there was moderate-to-high correlation between DEGs in excitatory neurons of PFC (Figure 4E), as well as between DEGs in inhibitory neurons of STR, at the two different ages (Figure 4F). Log_2_FCs of DEGs from non-neuronal cell types (astrocytes, microglia, and oligodendrocytes) also showed moderate-to-high correlation between 1- and 3-month-old *Sp4+/-* mutants (Figure 4G-L). Together, these data imply heterozygous LoF of *Sp4* causes large-scale, persistent changes in gene expression in both neuronal and non-neuronal cell types in PFC and STR.

### Cell type-specific dysregulations of diverse molecular pathways in *Sp4* mutants

The expression of activity-regulated genes can be used as a surrogate measure of neuronal activity in the brain^10,59^ (Table S1). Based on the GSEA of snRNA-seq data using the curated set of activity-regulated genes, we found that in *Sp4+/-* PFC, activity regulated genes were significantly enriched among the downregulated genes in L2/3IT neurons but among the upregulated genes in L5/6NP and Vip neurons at 1 month (Figure 5A; Figure S4A), suggesting reduced neuronal activity in L2/3IT and enhanced neuronal activity in L5/L6NP and Vip neurons at 1 month. More interestingly, at 3 months of age, activity-regulated genes were upregulated in essentially all neuronal cell types, both excitatory and inhibitory neurons, of PFC in *Sp4+/-* mutants (Figure 5A; Figure S4B). A similar pattern occurred in *Sp4+/-* STR, where the activity-regulated gene-set was downregulated in SPNs (the major inhibitory projection neuron of STR) at 1 month but became upregulated at 3 months, most significantly in direct pathway SPNs (dSPN that expresses *Drd1*) (Figure 5B-D). Thus, the transcriptomic data suggest that in *Sp4+/-* mutants, there is a juvenile period (∼1 month of age) of hypoactivity in the major neuronal cell types of PFC and STR, but this switches later to robust hyperactivity in early adulthood (3 months of age) (Figure 5A and B).

**Figure 5.**
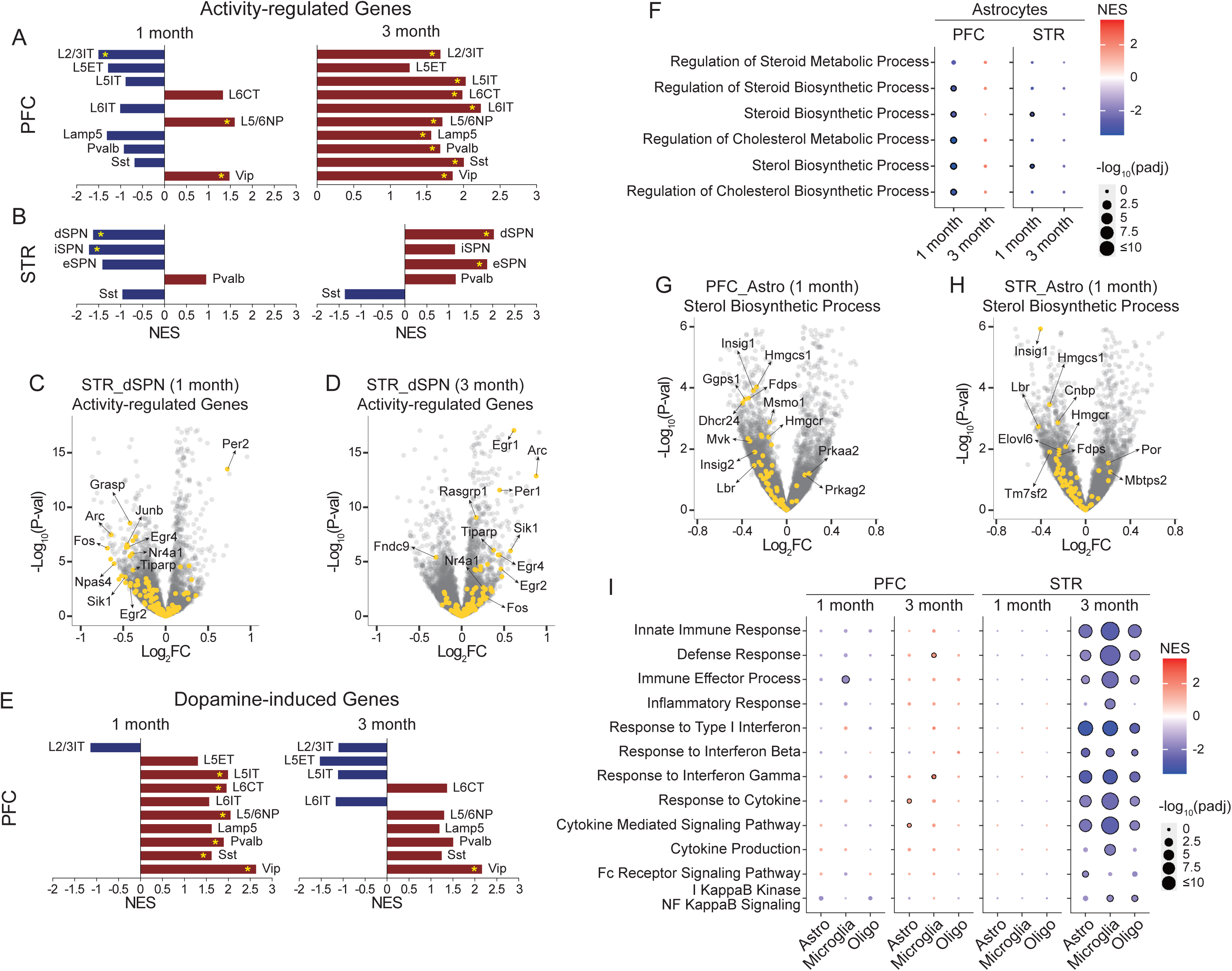
Cell type-specific effects on activity-, dopamine-, cholesterol-, and immune-related pathways in *Sp4+/−* mutant mice. (A and B) Enrichment of the activity-regulated gene set across neuronal subtypes in PFC (A) and STR (B) of *Sp4+/−* mutant mice at the indicated ages. (C and D) Volcano plots of transcriptomic changes in the indicated neuronal subtypes of STR in *Sp4+/-* mutant mice at 1 month (C) and 3 months (D), highlighting genes from the “Activity-regulated Genes” gene set. Log_2_FC, log_2_Fold Change. (E) Enrichment of the dopamine-induced gene set across neuronal subtypes in PFC of *Sp4+/−* mutant mice at the indicated ages. (F) GSEA results for the cholesterol-related GO terms in astrocytes of the indicated brain regions in *Sp4+/-* mutant mice at the indicated ages. (G and H) Volcano plots of transcriptomic changes in astrocytes of PFC (G) and STR (H) in 1-month-old *Sp4+/-* mutant mice, highlighting genes from the “Sterol Biosynthetic Process” GO term. Log_2_FC, log_2_Fold Change. (I) GSEA results for the immune-related GO terms in the indicated glial subtypes of PFC and STR in *Sp4+/-* mutant mice at the indicated ages. In (A), (B), and (E), yellow asterisks indicate statistical significance (adjusted p-value < 0.05). In (F) and (I), circles with black outlines indicate statistical significance (adjusted p-value < 0.05). NES, normalized enrichment score. See also Figure S4.

In the excitatory neurons of PFC of *Sp4+/-* mutants, genes associated with dopamine and serotonin GO terms (“Response to Dopamine”, “Response to Catecholamine”, “Serotonin Receptor Activity”, and “Positive Regulation of MAP Kinase Activity”), such as *Alk*, *Chrm2*, *Htr4*, and *Htr2c* were upregulated, particularly in L5IT or L5/6NP at 1 month of age (Figure S4C-E). This finding is similar to *Grin2a* mutant mice that also showed upregulated dopamine-related GO terms in excitatory neurons of PFC, particularly L5IT neurons^10^. Similar to PFC, the striatal inhibitory SPNs also showed a positive GSEA signal for “Response to Dopamine”, “Response to Catecholamine”, and “Serotonin Receptor Activity” GO terms (Figure S4C and F), although these did not reach significance, possibly due to the limited statistical power of our snRNA-seq study. Supporting the notion that there is elevated dopamine signaling in PFC neurons at 1 month, we found a significant enrichment of a set of dopamine-induced genes (Table S1) among the upregulated genes in L5IT, L6CT, L5/6NP, Pvalb, Sst, and Vip neurons of *Sp4+/-* PFC at 1 month; however, this GSEA signal weakened considerably at 3 months (Figure 5E; Figure S4G and H). Together, these results suggest that *Sp4* mutant mice have a hyperdopaminergic state particularly in PFC during the juvenile period.

Among non-neuronal cell types of *Sp4+/-* mutant mice, we found that genes related to cholesterol/steroid biosynthesis, including *Insig1*, *Hmgcs1*, *Hmgcr*, and *Lbr*, were significantly downregulated in astrocytes of PFC and STR at 1 month, but not at 3 months (Figure 5F-H). These findings are reminiscent of the recent findings from postmortem human SCZ brains showing reduced expression of synaptic neuron and astrocyte program (SNAP)^60^. In this regard, it is interesting that the downregulation of sterol biosynthesis genes at 1 month but not at 3 months is correlated with reduced activity-regulated genes in the cortical and striatal neurons in *Sp4+/-* mutants at 1 month (see Figure 5A and B).

A growing body of evidence has implicated immune dysfunction or neuroinflammation in the pathophysiology of SCZ and BD^60–63^. Notably, our GSEA of snRNA-seq data revealed downregulation of a range of immune-related GO terms (e.g. “Innate Immune Response”, “Response to Type I Interferon”, and “Response to Cytokine”) in glial cells of STR in particular (Figure 5I). This was found most strongly in microglia, but also in astrocytes and oligodendrocytes, and occurred at 3 months, but not at 1 month (Figure 5I; Figure S4I). Downregulated genes in STR microglia included *Irf1*, *Irf7*, *Ifitm3*, *Stat1*, and *Stat2* (Figure S4I). Unlike STR, the PFC glial cells showed minimal alteration in these immune-related GO terms at 1 and 3 months (Figure 5I). In addition, we found that genes associated with disease associated microglia (DAM)^61^ and homeostatic microglia^61^ were significantly enriched among down- and up-regulated genes, respectively, in the striatal microglia in *Sp4+/-* mutants at 1 and 3 months (Figure S4J-L). In PFC, genes associated with DAM and homeostatic microglia (Table S1) were significantly enriched among down-regulated genes at 1 month, but not at 3 months (Figure S4J). Moreover, genes associated with pan-reactive astrocytes and disease associated astrocytes (DAA)^62^ (Table S1) were significantly enriched among down-regulated genes in the striatal astrocytes and oligodendrocytes of 3-month-old *Sp4+/-* mutants (Figure S4J). Thus, our GSEA analysis indicates that microglia and astrocytes in *Sp4+/-* brain (most prominently in STR) change their “activation” state in essentially the opposite direction compared with microglia and astrocytes in neurodegenerative diseases^63–65^. Since *Sp4* is minimally expressed in glial cells, we speculate that gene expression changes in the observed pathways in astrocytes and microglia are secondary to the changes in neurons.

### Perturbed transcription factor gene expression programs in *Sp4+/-* mutant mice

*Sp4* is itself a TF and may regulate expression of other TFs that could lead to secondary effects on a wide range of gene-expression programs. To infer which TFs might account for the large transcriptomic changes found by snRNA-seq in various neuronal cell types of *Sp4+/-* mutant mice, we performed TF enrichment analysis based on orthogonal omics integration^66^ (see Methods). Among the TFs derived from the enrichment analysis, we focused on TFs that are themselves DEGs in each cell type of *Sp4+/-* mutants. This analysis revealed that 12 and 15 TFs accounted for 62.8% and 73.7% of the down- and up-regulated DEGs, respectively, in the cortical (PFC) L2/3IT neurons of 1-month-old *Sp4+/-* mutants (Figure 6A). In the cortical L5IT neurons of 1-month-old *Sp4+/-* mutants, 43.8% and 79.9% of the down- and up-regulated DEGs could be accounted for by 3 and 15 TFs, respectively (Figure 6B). Finally, 3 and 14 TFs were responsible for 56% and 85.3% of down- and up-regulated DEGs, respectively, in the cortical L6CT neurons of *Sp4+/-* mutants at 1 month (Figure 6C). It is interesting that *Nr3c1*, the glucocorticoid (stress hormone) receptor (GR) associated with psychosocial stress^67^, *Creb1*, cAMP responsive element binding protein associated with cognitive processes^68^, and *Mitf*, a regulator of neuronal activity and plasticity^69^, appear to regulate the largest proportion of DEGs (both down- and up-regulated DEGs) in L2/3IT, L5IT, and L6CT neurons, respectively (Figure 6A-C). In STR, the TF enrichment analysis showed that 12 and 15 TFs could accounted for 73.9% of the down-regulated DEGs in dSPNs and iSPNs at 1 month (Figure 6D and E). Notably, *Kdm5b*, a histone demethylase that is genetically associated with intellectual disability and psychiatric disorders^70,71^, was responsible for ∼41% of the down-regulated genes in both dSPN and iSPN neurons (Figure 6D and E). Among the TFs that might underlie the up-regulated DEGs in dSPNs and iSPNs, *Ahr*, a regulator of dendrite arborization and synaptic maturation that plays an important role in differentiation of GABAergic neurons^72^, and *Thrb*, a nuclear receptor for thyroid hormone that is critical for brain development, especially GABAergic neuron maturation^73^, were responsible for the most changes (19.7% and 22.3%, respectively) (Figure 6D and E).

**Figure 6.**
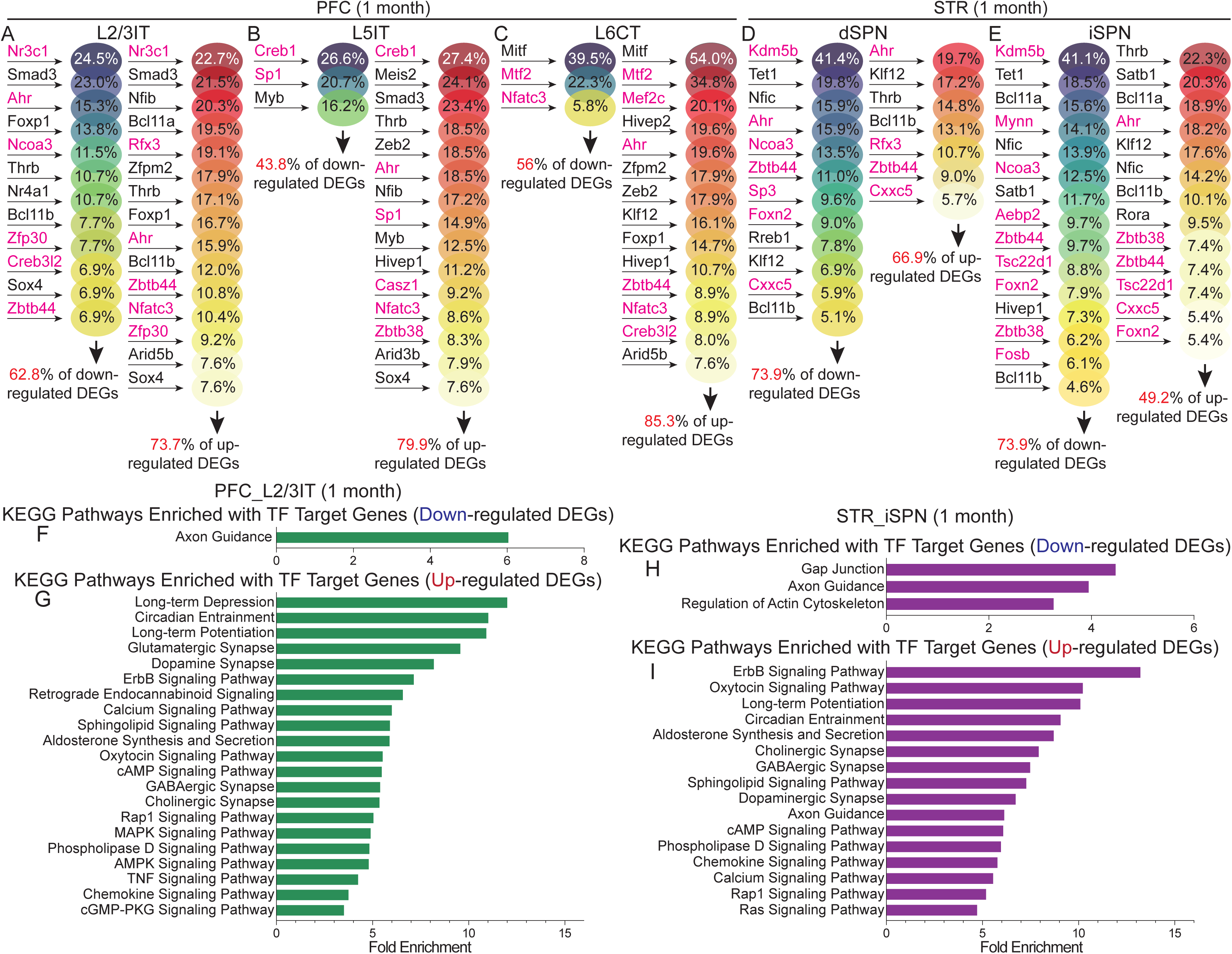
Dysregulated transcription factor gene expression programs in *Sp4+/-* mutant mice. (A-E) Transcription factor (TF) enrichment analysis results showing the TFs responsible for the down- or up-regulated DEGs and their percentage overlap between down- or up-regulated DEGs and target genes of the denoted TFs in the indicated neuronal subtypes of PFC (A-C) and STR (D and E) in 1-month-old *Sp4+/-* mutant mice. (F and G) KEGG pathway enrichment analysis of target genes of TFs responsible for the down-(F) or up-related (G) DEGs in L2/3IT of PFC in 1-month-old *Sp4+/-* mutant mice. (H and I) KEGG pathway enrichment analysis of target genes of TFs responsible for the down-(H) or up-related (I) DEGs in iSPN of STR in 1-month-old *Sp4+/-* mutant mice. In (A-E), the percentage of the total DEGs covered by the denoted TFs are written in red color below the black arrow. The magenta-colored genes are reported to be the direct targets of *Sp4*. See also Figure S5.

Although much less studied than *Sp1* and *Sp3*, potential target genes of *Sp4* have been recently identified by chromatin immunoprecipitation (ChIP)-seq experiments using postmortem human brains^74^. Notably, a high proportion (40-70%) of the TFs inferred from our TF enrichment analysis were *Sp4* target genes according to this ChIP-seq dataset (denoted in magenta color in Figure 6A-E). Those TFs included *Nr3c1*, *Kdm5b*, *Ahr*, *Ncoa3*, and *Mef2C*, which represent some of the top regulators of DEGs in the cortical excitatory and striatal inhibitory neurons at 1 month (Figure 6A-E). Overall, our analysis suggests that *Sp4* exerts a strong influence on neuronal gene expression in large part through regulation of a set of other TFs and chromatin modifiers.

To uncover the molecular pathways that are regulated by the target TFs of *Sp4* (such as *Nr3c1*, *Kdm5b*, *Creb1*, etc.), we performed over-representation analysis using ShinyGO^75^ on the target genes of these TFs obtained from the TF enrichment analysis. The target genes of the TFs that are responsible for the down-regulated DEGs in L2/3IT and L6CT of PFC and SPNs of STR were enriched in “Axon Guidance” KEGG pathway (Figure 6F and H; Figure S5B and D). The target genes of the TFs that are responsible for the up-regulated DEGs in L2/3IT, L5IT, and L6CT of PFC were enriched in KEGG pathways related to synapse (glutamatergic, dopaminergic, cholinergic, and GABAergic synapse), synaptic plasticity (long-term depression and long-term potentiation), and synaptic signaling (cAMP, MAPK, ErbB, and calcium signaling) (Figure 6G; Figure S5A and C). Similar KEGG pathways (LTP, dopaminergic synapse, cAMP signaling, etc.) were also regulated by the TFs responsible for up-regulated DEGs in SPNs of STR (Figure 6I; Figure S5E). It is interesting that KEGG pathways such as “Aldosterone Synthesis and Secretion”, “Oxytocin Signaling Pathway”, and “Thyroid Hormone Signaling Pathway” that are commonly altered in patients of psychiatric disorders including SCZ and BD^76–84^ were regulated by the driver TFs for the up-regulated DEGs in both the cortical and striatal neurons (Figure 6G and I; Figure S5A, C, and E). Collectively, these data imply that *Sp4* regulates a variety of TFs critical for brain function and neuronal signaling, thereby contributing to the pathophysiology of SCZ-BD psychotic disorders.

## Discussion

In this study, we studied the brains of *Sp4* mutant mice, a genetic model of SCZ and BD, using unbiased comprehensive multi-omics approaches to uncover molecular, cellular, and systems-level changes that might be relevant to disease pathophysiology. Here, we found that heterozygous LoF of *Sp4* results in prominent changes in gene expression across multiple brain regions, affecting similar genes and pathways as homozygous LoF of *Sp4* (including synapse, oxidative phosphorylation, and ribosome). According to our synapse proteomics data, various synaptic vesicle-related and glutamatergic signaling pathways are reduced at the synapses of *Sp4* mutant cortex, supporting the longstanding hypo-glutamate/NMDAR hypothesis of SCZ. In addition, *Sp4* LoF results in striking transcriptomic changes in both neuronal and non-neuronal cell types: dysregulations in activity-regulated gene expression and dopaminergic signaling in neurons, cholesterol biosynthesis in astrocytes, and immune response and inflammatory signaling in glial cells.

Furthermore, our study showed that the huge impact of *Sp4* LoF on neuronal gene expression could be due to largely through its regulation of a set of other TFs and chromatin modifiers that could lead to secondary effects on a wide range of gene-expression programs. Many of the TFs identified from our TF enrichment analysis are particularly interesting in SCZ and BD pathophysiology. For instance, *Sp4+/-* mutants showed decreased expression of *Nr3c1* and *Kdm5b* in neurons of PFC and STR at 1 month, respectively (data not shown). Consistently, reduced mRNA expression of GR (*Nr3c1*) has been found in hippocampus, dorsolateral prefrontal cortex, inferior temporal cortex, and amygdala in SCZ patients, and in the hippocampus, entorhinal cortex, inferior temporal cortex, and amygdala in BD patients^85,86^. Also, methylation of *Nr3c1* gene has been closely associated with psychosocial stress reactivity and hypothalamic–pituitary–adrenal (HPA) axis modulation^67^. Given that deregulated HPA axis has been reported in patients with various psychiatric disorders and people who experienced early psychosocial stress^67^, decreased expression of *Nr3c1* due to LoF of *Sp4* might contribute to pathophysiology of SCZ and BD.

In addition to *Kdm5b*, many other histone demethylases are the target genes of *Sp4* based on the recent human *Sp4* ChIP-seq data and significantly reduced in striatal neurons of *Sp4+/-* mutants at 1 month according to our snRNA-seq data (Figure S5F). Of note, among these histone demethylases, *Kdm6b* is also a rare-variant large-effect risk gene for SCZ identified by SCHEMA study. Previously, it has been suggested that *Sp4* might regulate the expression of many SCZ risk genes by directly binding to their GC-rich promotor regions^87^. Indeed, many SCZ risk genes identified from SCHEMA study were targets of human *Sp4* and were significantly altered in the neurons of STR in *Sp4+/-* mutants at 1 month (Figure S5G). Therefore, our study suggests that *Sp4* is not only an important risk gene for SCZ, but also serves as a central node in the network regulating other SCZ risk genes.

In addition to being a key intracellular effector of dopamine receptors, cAMP/PKA plays a key role in synaptic plasticity^88–91^. However, increased cAMP-PKA-calcium signaling has been reported in SCZ and BD patients^92,93^. Genetic evidence supports the involvement of PKA dysregulation in mood disorders, with the cAMP synthesizing enzyme Adcy2 being a major association in human GWAS studies for BD^94^. In addition, several mouse models of SCZ and BD, such as *Setd1a*, *Trio*, and *Akap11*, whose ultra-rare LoF variants are associated with greatly increased risk for SCZ and BD^8^, also showed up-regulation of cAMP-PKA pathway^95–97^. Since cAMP signaling was commonly enriched by the target genes of the TFs identified from our TF enrichment analysis in neuronal cell types of PFC and STR in *Sp4+/-* mutants at 1 month (see Figure 6 and Figure S5), future studies investigating whether *Sp4+/-* mutants also have dysregulated cAMP/PKA signaling might be very interesting. Overall, this study provides a molecular-to-systems-level picture of *Sp4* mutant brain and enables deeper understanding of SCZ and BD pathophysiology and disease mechanism.

## Methods

### Animals

*Sp4* mutant mice were generated using the CRISPR/Cas9 technique by introducing an early terminating stop codon in Exon3 (Figure 1A), where *Sp4* genetic variants are concentrated in SCZ patients. Sp4 protein levels in brains were validated to be reduced in heterozygous (*Sp4+/-*) and undetectable in homozygous (*Sp4-/-*) mutant mice compared to those of WT (*Sp4+/+*) littermates. Before used in various experiments described in this study, *Sp4* mutant mice were backcrossed against C57/BL6J mice (Jackson Laboratory, #000664) at least five generations. *Sp4+/-* mutant mice were crossed with *Sp4+/-* mutant mice to produce *Sp4+/+*, *Sp4+/-*, and *Sp4-/-* littermates used in bulk RNA-seq experiment. For snRNA-seq experiment, *Sp4+/-* mutant mice were crossed with their *Sp4+/+* littermates to produce *Sp4+/+* and *Sp4+/-* mice. To minimize variability, only 1- and 3-month-old *Sp4+/+*, *Sp4+/-*, and *Sp4-/-* male mice were used in this study. All animals were housed at AAALAC-approved facilities on a 12-hour light/dark cycle, with food and water available *ad libitum*. All procedures involving *Sp4* mutant mice were approved by the Broad Institute IACUC (Institutional Animal Care and Use Committee) and conducted in accordance with the NIH Guide for the Care and Use of Laboratory Animals.

### DNA isolation and genotyping

Tissues collected from ear punching were used to isolate DNAs as described in Jackson Laboratory protocol^98^ for genotyping. For genotyping, DNA oligo primers were ordered from IDT (Integrated DNA Technologies) with Standard Desalting purification. After DNA isolation, PCR was performed with KAPA HiFi HotStart ReadyMix PCR Kit (Roche, KK2601) following the manufacturer’s instructions, except that the extension time and the final extension time were set as 15 seconds and 1 minute, respectively. The PCR primers that specifically capture WT or mutated *Sp4* allele (Figure 1A and B) were as shown below:

Sp4 WT Forward Primer – CTAGAACTGGTGACGACGCA (20 bases)
Sp4 WT Reverse Primer – TAGCAGCCGTGAGGATAGCTT (21 bases)
Sp4 Mutant Forward Primer – CAGCTCATTTCTGCAGGTAATTTCG (25 bases)
Sp4 Mutant Reverse Primer – CACTGGGGTGATGGTTAACTGC (22 bases)

For these primers, the annealing temperature was set as 68°C, and the number of PCR cycles for WT and mutant mice were 35 and 40 cycles, respectively.

### Western Blot (WB)

To confirm Sp4 protein levels in the generated *Sp4* mutant mice, the brain tissues of these mice were mechanically homogenized with disposable pestles and cordless pestle motor in ice-cold RIPA buffer with protease inhibitor (Sigma, 4693159001) and phosphatase inhibitor (Sigma, 4906837001). The homogenate was centrifuged at 1,400g for 10 minutes, and the supernatant was collected. To measure protein concentration, bicinchoninic acid (BCA) protein assay (Thermo Fisher Scientific, A53225) was used, and the samples were prepared to be equal protein concentrations and volumes with the homogenization buffer. NuPAGE LDS Sample Buffer (4x, Thermo Fisher Scientific, NP0007) and NuPAGE Sample Reducing Agent (10x, Thermo Fisher Scientific, NP0009) were added to make the final WB loading samples. The samples were then heated at 95°C for 5 minutes before loading on to the Bolt 4-12% BisTris Gel (Thermo Fisher Scientific). After electrophoresis, the gel was transferred to a 0.2 µm Nitrocellulose membrane (Bio-Rad, Trans-Blot Turbo Mini 0.2 µm Nitrocellulose Transfer Packs, #1704158) for 30 minutes. For blocking, the membrane was incubated in the blocking buffer (5% skim milk in 0.1% TBST (Tris Buffered Saline + Tween 20) washing buffer) for 1 hour on the shaker at room temperature. After blocking, the membrane was incubated in the primary antibody solution (diluted in blocking buffer) on the shaker at 4°C for overnight. The secondary antibody incubation was performed for 1 hour. 0.1% TBST washing buffer was used for washing steps. 1:250 Mouse anti-Sp4 antibody (B-1, Santa Cruz Biotechnology, sc-390124) followed by 1:25,000 anti-mouse antibody (Thermo Fisher Scientific, # 31430), and 1:50,000 Mouse anti-β-actin HRP-conjugated antibody (MilliporeSigma, A3854) were used for this experiment.

### Brain perfusion and tissue collection for RNA sequencing

Brain tissues for bulk and snRNA-seq were collected as described in the previous study^10^ and protocols.io (https://www.protocols.io/view/fresh-frozenmouse-brain-preparation-for-single-nu-bcbrism6). Mice were anesthetized with isoflurane administration in a gas chamber. While anesthesia was prolonged via a nose cone through which 3% isoflurane flowed, transcardial perfusions were performed with ice-cold Hank’s Balanced Salt Solution (HBSS, Gibco, 14175-095) to remove blood from the brain. Immediately after perfusion, the brains were frozen in liquid nitrogen vapor and stored at –80°C until dissection.

Sample sizes used for bulk RNA-seq experiments were as shown below:

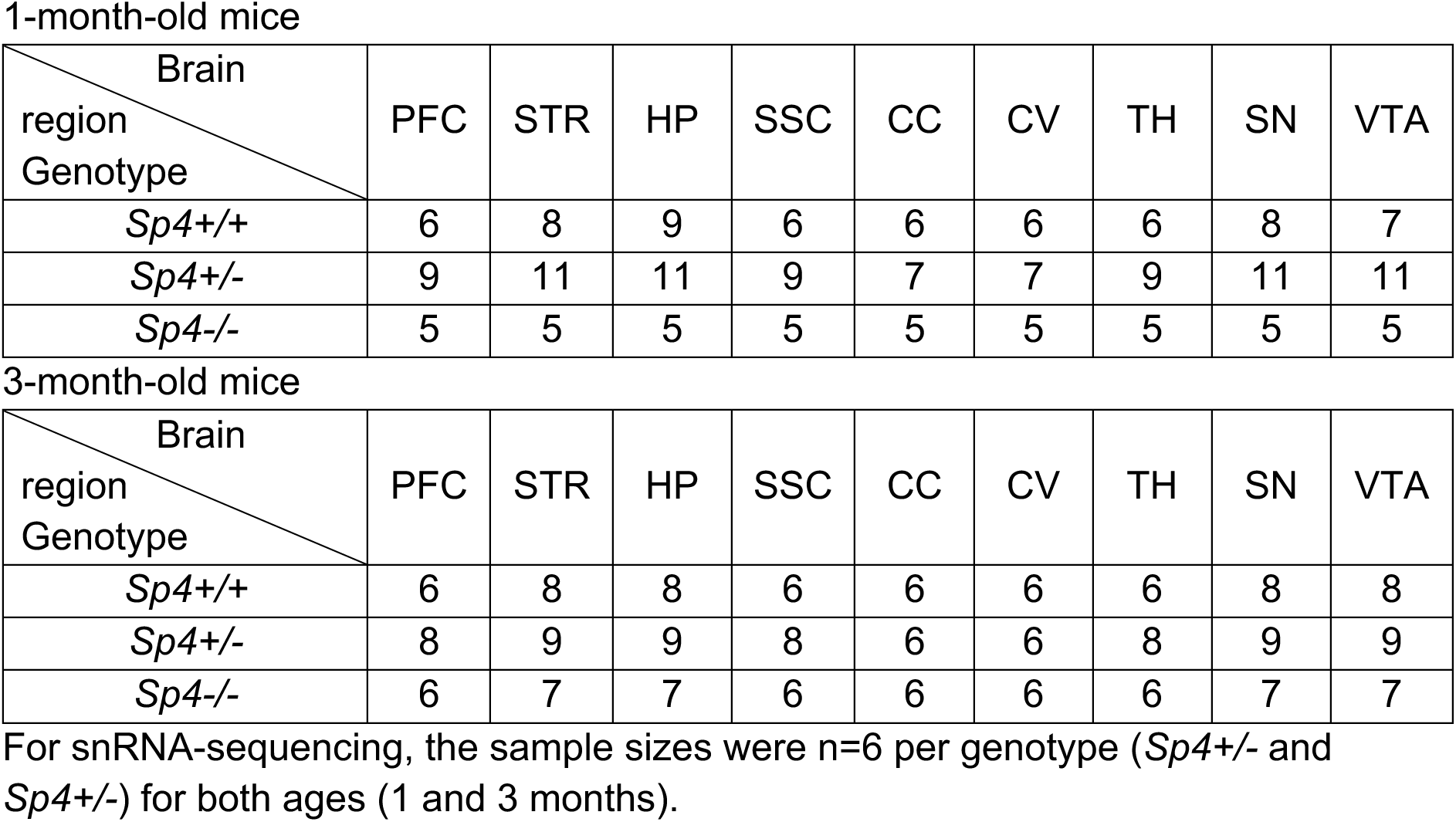

| Brain<br>region<br>Genotype | PFC | STR | HP | SSC | CC | CV | TH | SN | VTA |
| --- | --- | --- | --- | --- | --- | --- | --- | --- | --- |
| *Sp4*<sup>+/+</sup> | 6 | 8 | 9 | 6 | 6 | 6 | 6 | 8 | 7 |
| *Sp4*<sup>+/-</sup> | 9 | 11 | 11 | 9 | 7 | 7 | 9 | 11 | 11 |
| *Sp4*<sup>-/-</sup> | 5 | 5 | 5 | 5 | 5 | 5 | 5 | 5 | 5 |

3-month-old mice
| Brain<br>region<br>Genotype | PFC | STR | HP | SSC | CC | CV | TH | SN | VTA |
| --- | --- | --- | --- | --- | --- | --- | --- | --- | --- |
| *Sp4*<sup>+/+</sup> | 6 | 8 | 8 | 6 | 6 | 6 | 6 | 8 | 8 |
| *Sp4*<sup>+/-</sup> | 8 | 9 | 9 | 8 | 6 | 6 | 8 | 9 | 9 |
| *Sp4*<sup>-/-</sup> | 6 | 7 | 7 | 6 | 6 | 6 | 6 | 7 | 7 |

For snRNA-sequencing, the sample sizes were n=6 per genotype (*Sp4*<sup>+/-</sup> and *Sp4*<sup>-/-</sup>) for both ages (1 and 3 months).

Brain region microdissections were performed by hand in a cryostat (Leica CM3050S) with an ophthalmic microscalpel (Feather safety Razor no. P-715), 1.0 mm diameter biopsy punch (Thermo Fisher Scientific, Integra 33-31A), or 1.5 mm diameter biopsy punch (Thermo Fisher Scientific, Integra 33-31A). The medial PFC, HP, and SSC were dissected by hand in the cryostat using a precooled (to –20°C) ophthalmic microscalpel (Feather safety Razor no. P-715). The dorsal STR and TH were collected by a precooled 1.5 mm diameter biopsy punch while a precooled 1.0 mm diameter biopsy punch was used to collect SN and VTA. All brain regions were identified according to the Allen Brain Atlas. Dissected tissues were placed into a precooled 1.5-ml PCR tube and stored at –80°C until RNA extraction.

### RNA extraction and bulk RNA-seq library preparation

RNAs from the micro-dissected tissues were extracted using RNeasy Mini Kit (Qiagen) following the manufacturer’s instructions. RNA concentration was measured using a NanoDrop Spectrophotometer and RNA integrity (RIN) was measured with RNA pico chips (Agilent) using a 2100 Bioanalyzer Instrument (Agilent). Purified RNA was stored at -80°C until bulk RNA-seq library preparation. Bulk sequencing libraries were prepared using TruSeq Stranded mRNA Kit (Illumina) following the manufacturer’s instructions. 200 ng of isolated total RNA from each sample was used and the concentration of resulting cDNA library was measured with High Sensitivity DNA chips (Agilent) using a 2100 Bioanalyzer Instrument (Agilent). A 10 nM normalized library was pooled, and sequencing was performed on NovaSeq S2 (Illumina) with 50 bases each for reads 1 and 2 and 8 bases each for index reads 1 and 2.

### Nuclei extraction and snRNA-seq library preparation

Brain tissue dissection and nuclei extraction were performed within 24 hours to avoid desiccation and freeze-thaw cycles. A gentle, detergent-based dissociation was used to extract the nuclei according to a previously published protocol^10,99^, also available at protocols.io (https://www.protocols.io/view/frozen-tissue-nuclei-extraction-bbseinbe). Extracted nuclei were then loaded into the 10x Chromium V3.1 system (10x Genomics) and library preparation was performed following the manufacturer’s instructions. A 10 nM normalized library was pooled, and sequencing was performed on NovaSeq S2 (Illumina) with 28 and 75 bases for reads 1 and 2 and 10 bases each for index reads 1 and 2.

### Bulk RNA-seq data processing

Raw FASTQ files were obtained from each sequencing experiment and re-aligned to a standardized reference (genome FASTA file and transcriptome GTF were extracted from the CellRanger mm10 reference and used for alignment and quantification). Quantification was performed by Salmon^100^ (version 1.7.0, with arguments -l A --posBias --seqBias --gcBias --validateMapping), using a Salmon reference built with the salmon index command with genomic decoys included. Quality metrics were obtained from STAR alignment^101^ (version 2.7.10a) and Picard tools (https://broadinstitute.github.io/picard/, version 2.26.7). Sample P30_SN_WT_8 was removed from analysis for low read depth and poor library QC. Samples 3M_STR_HT_8 and 3M_TH_HT_3 were removed due to gene expression indicative of misdissection/contamination.

### Bulk RNA-seq differential expression (DE) analysis

DE analysis was performed between heterozygous mutants and WT mice for each individual experiment (age and brain region) using R package DESeq2^102^ (version 1.34). Salmon output was loaded using R package tximport^103^ (version 1.22). Log_2_ fold change shrinkage was applied using the “normal” shrinkage estimator in DESeq2. Differentially expressed genes (DEGs) were defined as genes with padj < 0.05.

For comparing the transcriptomic changes in *Sp4+/-* versus *Sp4-/-*, the Spearman’s correlation r values were calculated between Log_2_FC values of the same brain regions in *Sp4+/-* versus *Sp4-/-*. It should be noted that the use of the same set of WTs for *Sp4+/-* and *Sp4-/-* DE analysis may lead to over inflated correlation estimates. GSEA was performed with the R Bioconductor package fgsea^104^ using C5 ontology gene sets obtained from the Molecular Signatures Database (MSigDB) and SynGO (version 1.2).

### snRNA-seq data processing

FASTQs were generated from raw BCL files then aligned to a mm10 mouse reference genome using the Cell Ranger pipeline^105^ (v6.1.2). For Cell Ranger count, --chemistry=SC3Pv3 was used and --expect-cells=15,000. Within each experiment (i.e. brain region and age), replicates were down-sampled using Cell Ranger aggr. Seurat^106^ (v4.0.3) was used to further process and analyze the Unique Molecular Identifier (UMI) counts. Nuclei expressing less than 500 genes were removed. The data from remaining nuclei was log-normalized and scaled by a factor of 10000, then scaled with ScaleData to prepare for dimension reduction. Linear dimension reduction was performed by RunPCA on variable genes. The nuclei were then clustered using FindNeighbors (with 20 dimensions) and FindClusters and visualized using Uniform Manifold Approximation and Projection (UMAP) using 20 dimensions. Doublet identification was performed by Scrublet^107^ (v0.2.3), and nuclei above a threshold determined by the score distribution were removed, as well as small clusters that were labeled in majority as doublets. Major cell types were labeled by examining expression of marker genes, and neuronal nuclei were re-clustered, and subtypes were labeled using Azimuth^108^ (v0.4.6). Cell type proportions were summarized, and differences were statistically evaluated using speckle’s^104^ (v0.0.3) propeller function. GSEA was performed with the R Bioconductor package fgsea^104^ using C5 ontology gene sets obtained from the Molecular Signatures Database (MSigDB).

### snRNA-seq DE analysis

For DE analysis, a pseudocell approach^58^ was employed for each age and brain region. We utilized the randomized pseudocell generation approach, which aggregates expression of every 20-40 nuclei with similar transcriptomes within each replicate and cell type or subtype grouping. Cell types or subtypes that had too few nuclei to generate at least two pseudocells per replicate were removed. Limma’s duplicate Correlation mixed model analysis function was used to perform differential expression analysis with robust empirical Bayes moderated t-statistics and mouse ID as random effect^109^. Percentage of mitochondrial reads and genotype were used as covariates for all DE analyses. DEGs were defined as genes with Benjamini-Hochberg adjusted p-value < 0.05. Robustness of the identified DEGs was tested by calculating the log fold-change pattern (summarized as up or down regulated) of pairwise combinations of *Sp4+/-* versus *Sp4+/+* samples. We then calculated the fraction of comparisons for which the Log_2_FC had a similar direction of effect to the mixed linear model. We further filter DEGs to those which have an absolute robustness score > 0.6. GSEA was performed with the R Bioconductor package fgsea^104^ using C5 ontology gene sets obtained from the MSigDB.

### TRADE

For each brain region at each age, separately for *Sp4+/-* and *Sp4-/-* mutants, we estimated differential expression effects using DESeq2. We included surrogate variables estimated with SVA^110^ as fixed effects in the DESeq2 model. We picked the number of surrogate variables using the permutation-based ’be’ method included in SVA. We then submitted the DESeq2 summary statistics for analysis with TRADE^27^ to estimate the distribution of differential expression effects. We extracted the variance of the effect size distribution to determine the “transcriptome-wide impact” of *Sp4* mutation in each brain region at each age. To assess significance of these estimates, we repeated the above analysis 1000 times with permuted genotype labels and computed the proportion of permutation transcriptome-wide impact estimates that were larger than the empirical estimate.

### Transcription Factor Enrichment Analysis

For transcription factor enrichment analysis, the enrichment of up- and down-regulated DEGs was tested using ChEA3’s overrepresentation test^66^ for each cell type. We selected TFs of interest by identifying those which were differentially expressed in *Sp4+/-* mutants in the cell type of interest, were associated with Sp4 in the ChEA3 database, and showed FDR-significant enrichment in at least one of the ChEA3 libraries.

For each of these selected TFs, the percentage of DEGs calculated by counting the number of DEGs associated with that TF in any of the ChEA3 libraries (regardless of whether enrichment in that database was significant or not) and subsequently dividing that number by the total number of DEGs tested in the enrichment analysis. This was performed separately for up- and down-regulated DEGs.

To identify genes bound directly by SP4 in the brain, we obtained human SP4 binding sites from previously published ChIP-seq data from human DLPFC and cerebellum^74^. We obtained promoter site annotations in the human genome from the Eukaryotic Promoter Database^111^, then expanded these single base coordinates by 500bp upstream and 100bp downstream to approximate the promoter region. We then identified mouse genes whose promoter coordinates overlapped with human SP4 binding sites derived from the ChIP-seq and determined them as potential direct target genes of Sp4. For this comparison, we did human to mouse gene conversion since the ChIP-seq data is from human.

### Purification of synaptic fraction for mass spectrometry

1- and 3-month-old WT and *Sp4+/-* mutant mice were sacrificed with CO_2_ anesthesia, and their cortices were dissected rapidly, flash-frozen on dry ice, and stored at -80°C. Synapse fractions were purified as we described previously^36,112,113^. Briefly, flash-frozen cortices were homogenized by a teflon homogenizer and glass vessel in ice-cold homogenization buffer (5 mM HEPES pH 7.4, 1 mM MgCl2, 0.5 mM CaCl2, supplemented with phosphatase inhibitor (Sigma, 4906837001) and protease (Sigma, 4693159001) inhibitor. The homogenate was centrifuged for 10 minutes at 1,400 g 4°C, and the resulting supernatant (S1) was re-centrifuged at 13,800 g 4°C for 10 minutes. The resulting pellet (P2) was resuspended in 0.32 M Sucrose, 6 mM Tris-HCl (pH 7.5) and layered gently on a 0.85 M, 1 M, 1.2 M discontinuous sucrose gradient (all layers in 6 mM Tris-HCl pH 8.0) and ultracentrifuged at 82,500 g 4°C for 2 hours. The synaptosome fraction, which sediments at the interface between 1 M and 1.2 M sucrose, was collected. An equal volume of ice-cold 1% Triton X-100 (in 6 mM Tris-HCl pH 7.5) was added, mixed thoroughly, and incubated on ice for 15 minutes. The mixture was ultracentrifuged at 32,800 g 4°C for 20 minutes, and the pellet (the synapse fraction) was collected by resuspension in 1% SDS. A small aliquot was taken to measure the protein concentration using the Pierce Micro BCA assay (Thermo Fisher Scientific, 23235) and the remaining protein was stored at -80°C until being processed for MS. For synapse proteomics, seventeen synaptic fractions from WT and *Sp4+/-* mouse cortices were analyzed using tandem mass tag (TMT) isobaric labeling for quantification. Sample sizes were n=4 for both WT and *Sp4+/-* at 1 month, and n=4 for WT and n=5 for *Sp4+/-* at 3 months.

### Proteomics analysis of synapse fractions

The protein in synapse fraction samples (in 1% SDS) was reduced using 5 mM dithiothreitol and alkylated using 10 mM iodoacetamide at room temperature. The denatured, reduced, alkylated protein samples were then processed using S-Trap sample processing technology (Protifi) following manufacturer’s instructions. The proteins were bound to the S-Trap column via centrifugation and contaminants/detergents were washed away. Sequential digestion steps were then performed on column using 1:20 enzyme to substrate ratio of Lys-C for 2 hours and Trypsin overnight at room temperature.

Following digestion and desalting, 13 μg of each sample was labeled with TMT18 reagent; the TMT18 plex was constructed by randomly assigning the samples from each group to channels within the plex. After verifying successful labeling of more than 99% label incorporation, reactions were quenched using 5% hydroxylamine and pooled. The TMT18 labeled peptides were desalted on a 50 mg tC18 SepPak cartridge and fractionated by high pH reversed-phase chromatography on a 2.1 mm x 250 mm Zorbax 300 extend-c18 column (Agilent). One-minute fractions were collected during the entire elution and fractions were concatenated into 12 fractions for LC-MS/MS analysis.

One microgram of each proteome fraction was analyzed on a Exploris 480 QE mass spectrometer (Thermo Fisher Scientific) coupled to a Vanquish Neo LC system (Thermo Fisher Scientific). Samples were separated using 0.1% Formic acid / Acetonitrile as buffer A and 0.1% Formic acid / Acetonitrile as buffer B on a 27cm 75um ID picofrit column packed in-house with Reprosil C18-AQ 1.9 mm beads (Dr Maisch GmbH) with a 90 minutes gradient consisting of 1.8-5.4% B in 1 minute, 5.4-27% B for 84 minutes, 27-54% B in 9 minutes, 54-81% B for 1 minute followed by a hold at 81% B for 5 minute. The MS method consisted of a full MS scan at 60,000 resolution and a normalized AGC target of 300% and maximum inject time of 10 ms from 350-1800 m/z followed by MS2 scans collected at 45,000 resolution with a normalized AGC target of 30% with a maximum injection time of 105 ms and a dynamic exclusion of 15 seconds. Precursor fit filter was used with a threshold set to 50% and window of 1.2 m/z. The isolation window used for MS2 acquisition was 0.7 m/z and 20 most abundant precursor ions were fragmented with a normalized collision energy (NCE) of 34 optimized for TMT18 data collection.

### Proteomics data analysis

Mass spectra were analyzed using Spectrum Mill MS Proteomics Software (Broad Institute) with a mouse database from Uniprot.org downloaded on 04/07/2021 containing 55734 entries. Search parameters included: ESI Q Exactive HCD v4-35-20 scoring, parent and fragment mass tolerance of 20 ppm, 40% minimum matched peak intensity, trypsin allow P enzyme specificity with up to four missed cleavages and calculate reversed database scores enabled. Fixed modifications were carbamidomethylation at cysteine. TMT labeling was required at lysine, but peptide N termini could be labeled or unlabeled. Allowed variable modifications were protein N-terminal acetylation, oxidized methionine, pyroglutamic acid, and pyro carbamidomethyl cysteine. Protein quantification was achieved by taking the ratio of TMT reporter ions for each sample over the TMT reporter ion for the median of all channels. TMT18 reporter ion intensities were corrected for isotopic impurities in the Spectrum Mill protein/peptide summary module using the afRICA correction method which implements determinant calculations according to Cramer’s Rule and correction factors obtained from the reagent manufacturer’s certificate of analysis (https://www.thermofisher.com/order/catalog/product/90406) for lot numbers WG333575 and XC342532. After performing median normalization, a moderated two-sample t-test was applied to the datasets to compare WT and *Sp4+/-* sample groups at 1- and 3-months of age. Significant differentially expressed proteins (DEPs) were defined as proteins with nominal p-value < 0.01.

---

**Figure S1.**
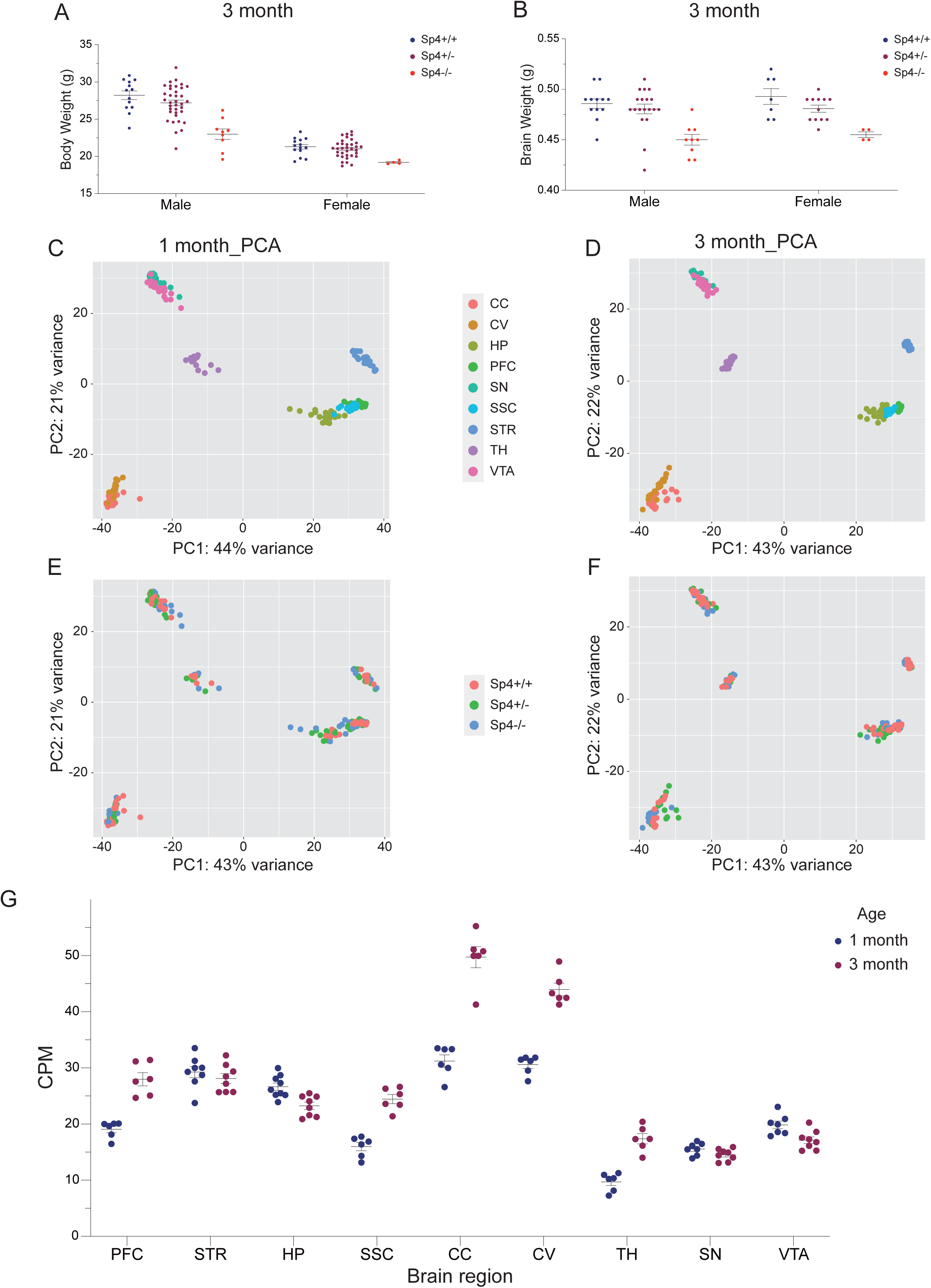
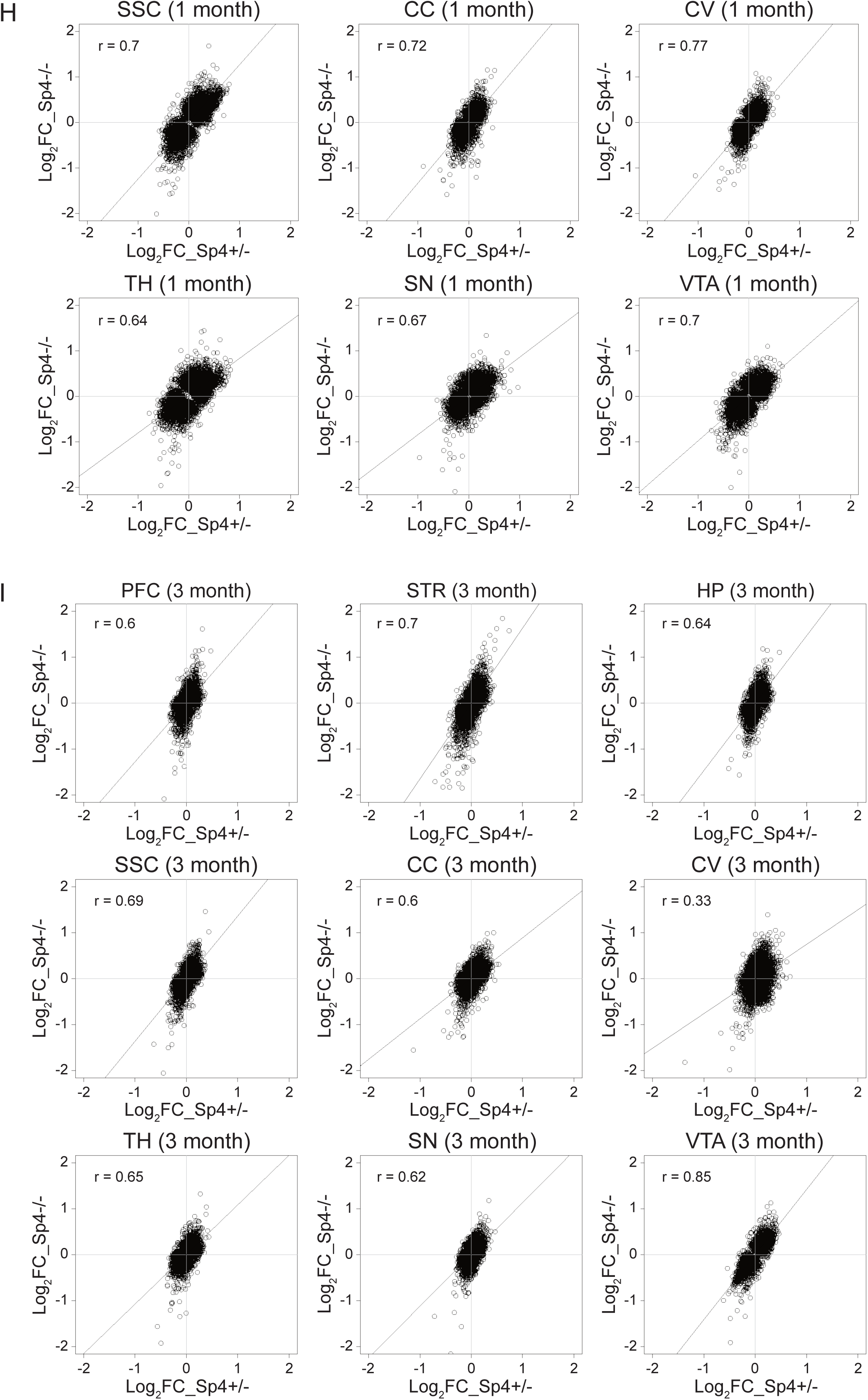
Brain-wide changes of transcriptome in *Sp4* mutant mice, Related to Figure 1. (A and B) Body (A) and brain (B) weights of the 3-months-old *Sp4+/-* and *Sp4-/-* mutants compared to WT littermates. (C-F) Principal component analysis (PCA) plot indicating the clustering of genotypes for the indicated brain regions from 1- and 3-month-old *Sp4* mutant mice and WT littermates. (G) *Sp4* expression in WT mice in the indicated brain regions and ages. CPM, counts per million. (H) Gene expression correlation in the indicated brain regions and genotypes at 1 month. Spearman’s r correlation values are indicated on the plots. Log_2_FC, log_2_Fold Change. (I) Gene expression correlation in the indicated brain regions and genotypes at 3 months. Spearman’s r correlation values are indicated on the plots. Log_2_FC, log_2_Fold Change. In (A-G), each circle represents an individual animal.

**Figure S2.**
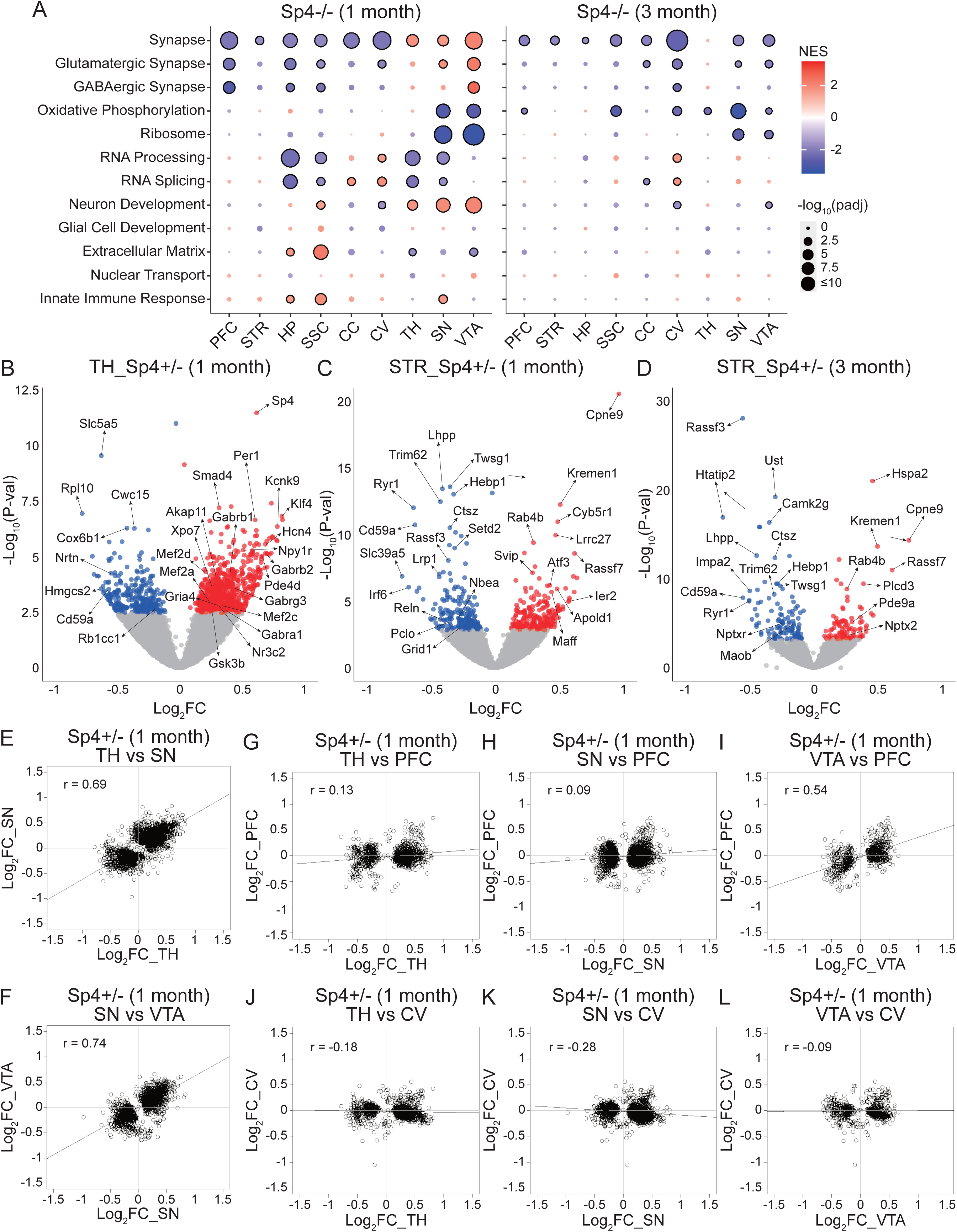
Transcriptomic changes in *Sp4* mutant mice, Related to Figure 2. (A) GSEA results of *Sp4-/-* mutants in the indicated brain regions and ages for a selection of GO terms from MSigDB. (B-D) Volcano plots of transcriptomic changes found in the bulk RNA-seq analysis of the indicated brain regions of *Sp4+/-* mutants (blue and red dots represent down- and up-regulated DEGs, respectively). Log_2_FC, log_2_Fold Change. (E-L) Correlation plots comparing the Log_2_FC of the DEGs between the denoted brain regions of *Sp4+/-* mutants at 1 month. In (A) and (B), circles with black outlines indicate statistical significance (adjusted p-value < 0.05). NES, normalized enrichment score. In (E-L), Spearman’s r correlation values are indicated on the plots. Log_2_FC, log_2_Fold Change.

**Figure S3.**
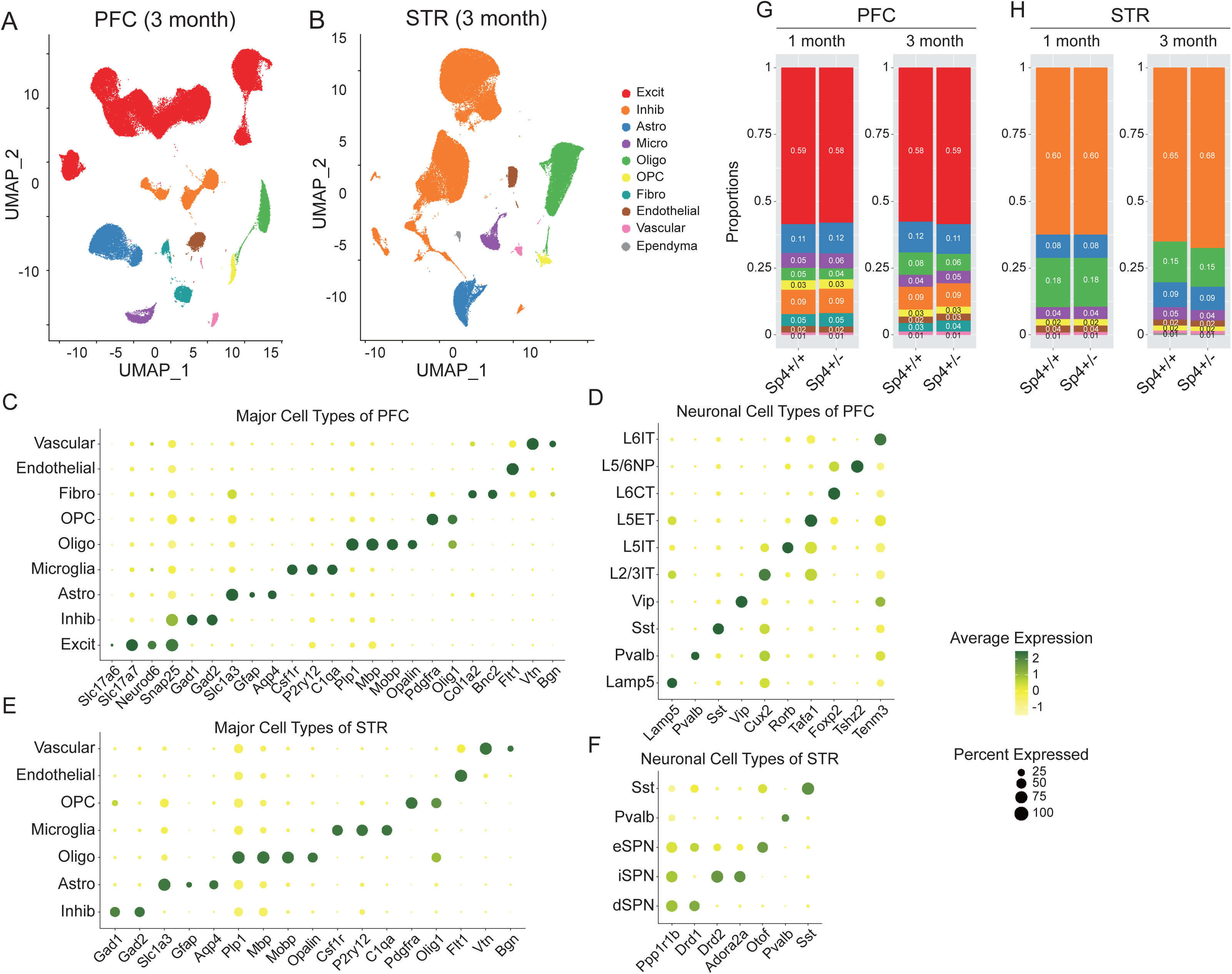
Brain region-specific cell types identified by snRNA-seq analysis in *Sp4* mutant mice, Related to Figure 4. (A and B) UMAP representation of the major cell types identified by snRNA-seq in the indicated brain regions at 3 months. (C-F) Dot plots describing the average and percentage of cells expressing cell type-specific marker genes in PFC (C and D) and STR (E and F) in 1-month-old *Sp4* animals. Similar marker genes were used to identify cell types in 3-month-old *Sp4* datasets. (G and H) Fraction of cell types identified in PFC (G) and STR (H) of animals at the indicated ages. Each column represents the average values for one genotype (N = 6 animals per genotype). No significant difference was observed in the proportion of identified cell types between *Sp4* mutant mice and their WT littermates.

**Figure S4.**
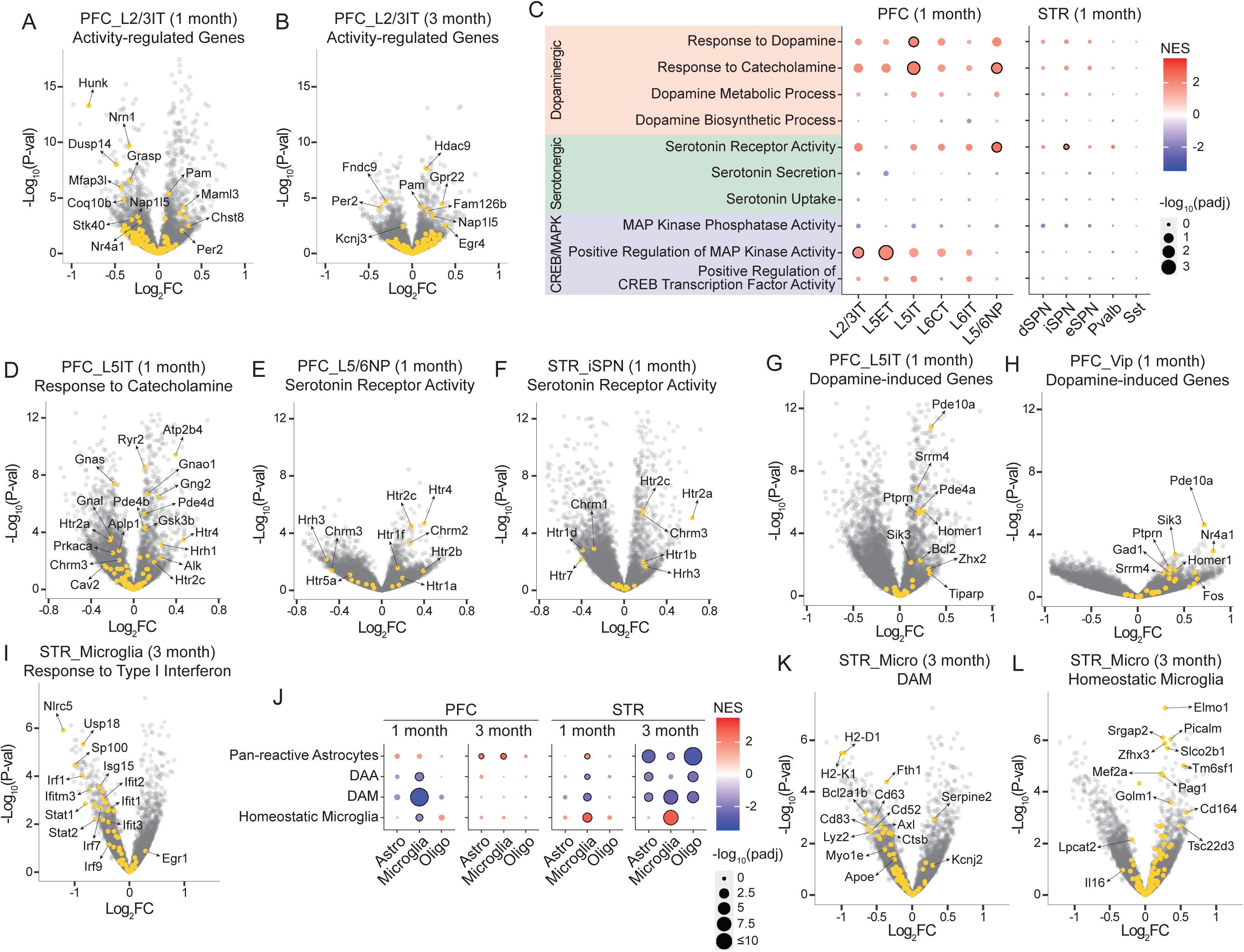
Brain-wide effects on activity-, dopamine-, and immune-related pathways across different cell types of *Sp4+/−* mutant mice, Related to Figure 5. (A and B) Volcano plots of transcriptomic changes in the indicated neuronal subtypes of PFC in *Sp4+/-* mutant mice at 1 month (A) and 3 months (B), highlighting genes from the “Activity-regulated Genes” gene set. Log_2_FC, log_2_Fold Change. (C) GSEA results for the indicated GO terms across neuronal subtypes of PFC and STR in 1-month-old *Sp4+/-* mutant mice. (D-F) Volcano plots of transcriptomic changes in the indicated neuronal subtypes of PFC (D and E) and STR (F) in 1-month-old *Sp4+/-* mutant mice, highlighting genes from the “Response to Catecholamine” (D) and “Serotonin Receptor Activity” (E and F) GO terms. Log_2_FC, log_2_Fold Change. (G and H) Volcano plots of transcriptomic changes in the indicated neuronal subtypes of PFC in 1-month-old *Sp4+/-* mutant mice, highlighting genes from the “Dopamine-induced Genes” gene set. Log_2_FC, log_2_Fold Change. (I) Volcano plot of transcriptomic changes in microglia of STR in 3-month-old *Sp4+/-* mutant mice, highlighting genes from the “Response to Type I Interferon” GO term. Log_2_FC, log_2_Fold Change. (J) Enrichment of the indicated gene sets in the glial subtypes of PFC and STR *Sp4+/-* mutant mice at the indicated ages. (K and L) Volcano plots of transcriptomic changes in the striatal microglia in 3-month-old *Sp4+/-* mutant mice, highlighting genes from the “DAM” (K) and “Homeostatic Microglia” (L) gene sets. Log_2_FC, log_2_Fold Change. In (C) and (J), circles with black outlines indicate statistical significance (adjusted p-value < 0.05). NES, normalized enrichment score.

**Figure S5.**
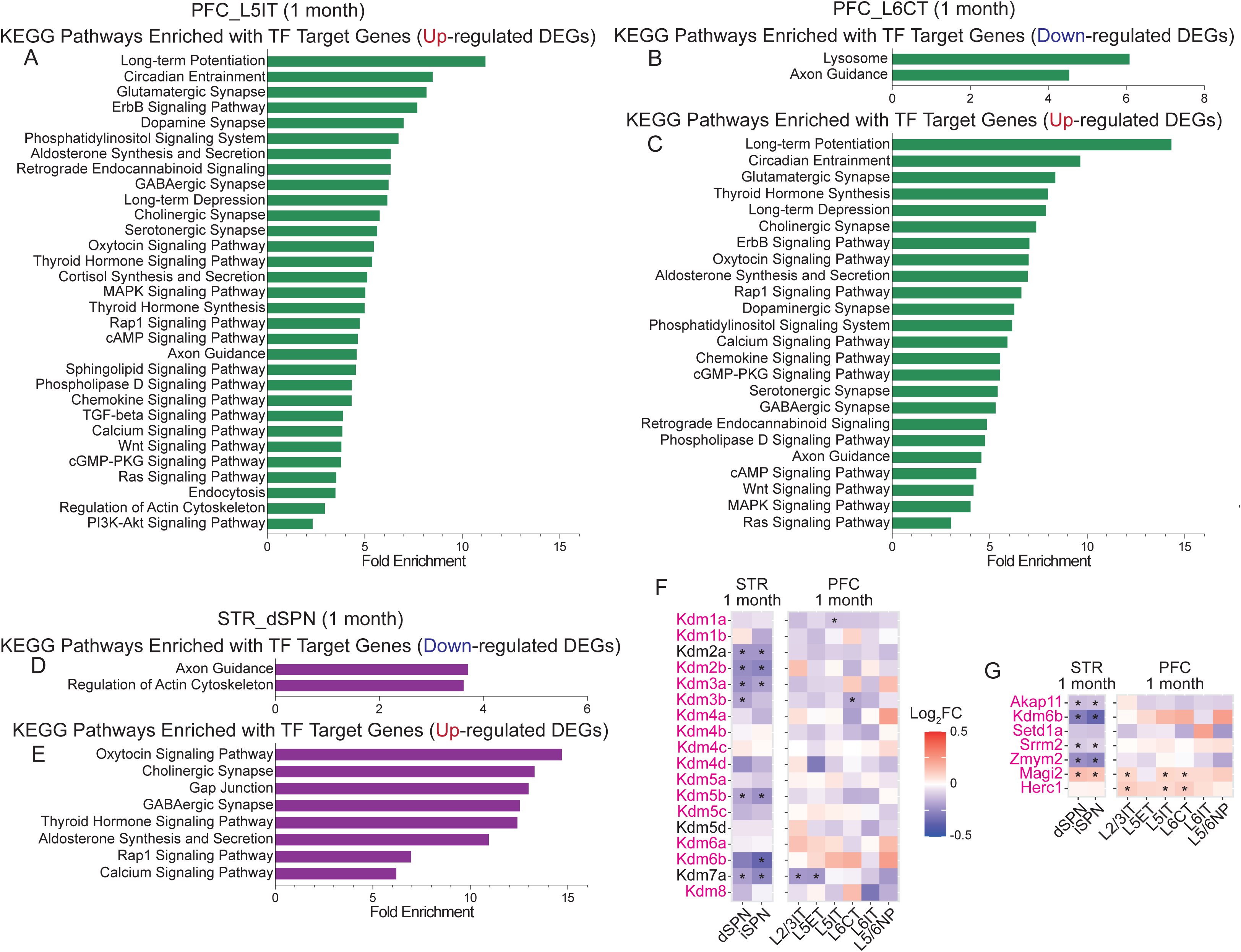
Perturbed gene expression programs resulting from LoF of *Sp4*, Related to Figure 6. (A) KEGG pathway enrichment analysis of target genes of TFs responsible for the up-related DEGs in L5IT of PFC in 1-month-old *Sp4+/-* mutant mice. (B and C) KEGG pathway enrichment analysis of target genes of TFs responsible for the down-(B) or up-related (C) DEGs in L6CT of PFC in 1-month-old *Sp4+/-* mutant mice. (D and E) KEGG pathway enrichment analysis of target genes of TFs responsible for the down-(D) or up-related (E) DEGs in dSPN of STR in 1-month-old *Sp4+/-* mutant mice. (F) Heatmap of gene expression changes of histone lysine demethylase genes across neuronal subtypes of PFC and STR in 1-month-old *Sp4+/-* mutant mice. (G) Heatmap of gene expression changes of SCHEMA genes that are reported to be the direct targets of Sp4 across neuronal subtypes of PFC and STR in 1-month-old *Sp4+/-* mutant mice. In (F) and (G), the magenta-colored genes are reported to be the direct targets of *Sp4*, and black asterisks indicate statistical significance (adjusted p-value < 0.05). Log_2_FC, log_2_Fold Change.

## Notes

### Competing Interest Statement

The authors have declared no competing interest.

